# Detection and Degradation of Nonsense-mediated mRNA Decay Substrates Involve Two Distinct Upf1-bound Complexes

**DOI:** 10.1101/266833

**Authors:** Marine Dehecq, Laurence Decourty, Abdelkader Namane, Caroline Proux, Joanne Kanaan, Hervé Le Hir, Alain Jacquier, Cosmin Saveanu

## Abstract

Nonsense-mediated mRNA decay (NMD) is a translation-dependent RNA degradation pathway involved in many cellular pathways and crucial for telomere maintenance and embryo development. Core NMD factors Upf1, Upf2 and Upf3 are conserved from yeast to mammals, but a universal NMD model is lacking. We used affinity purification coupled with mass spectrometry and an improved data analysis protocol to obtain the first large-scale quantitative characterization of yeast NMD complexes in yeast (112 experiments). Unexpectedly, we identified two distinct complexes associated with Upf1: Detector (Upf1/2/3) and Effector. Effector contained the mRNA decapping enzyme, together with Nmd4 and Ebs1, two proteins that globally affected NMD and were critical for RNA degradation mediated by the Upf1 C-terminal helicase region. The fact that Nmd4 association to RNA was dependent on Detector components and the similarity between Nmd4/Ebs1 and mammalian Smg5-7 proteins suggest that in all eukaryotes NMD operates through successive Upf1-bound Detector and Effector complexes.

## Introduction

Nonsense-mediated mRNA decay (NMD) is a major surveillance pathway defined by its importance for the fast decay of mRNAs with premature termination codons (PTC). Rapid translation-dependent RNA degradation prevents the production of truncated proteins (Hall & Thein, 1994) and reduces the loss of ribosomes “locked” on RNA at aberrant termination sites (Ghosh *et al*, 2010; Serdar *et al*, 2016). NMD factors are essential for the development of metazoans (Medghalchi *et al*, 2001) and participate to telomere maintenance and telomerase activity regulation in yeast (Addinall *et al*, 2011) and human cells (Azzalin *et al*, 2007). As a quality control pathway, NMD shapes the transcriptome of eukaryotic cells, both under normal and pathologic conditions, including cancer (Lindeboom *et al*, 2016). While NMD substrates were initially thought to be rather rare, single-nucleotide resolution RNA sequencing in yeast revealed that genome-wide transcription generates thousands of RNAs that are degraded by NMD (Malabat *et al*, 2015).

Large-scale results confirmed and validated previous studies on individual NMD reporters that described two classes of NMD substrates: one in which a long 3’ untranslated regions (UTR) extensions follows a short coding sequence and another that includes an exon-exon junction, bound by the exon-exon junction complex (EJC), downstream a stop codon. While long 3’-UTRs destabilize RNAs in all the eukaryotes studied to date, EJC-enhanced NMD is absent from *S. cerevisiae*, *S. pombe* (Wen & Brogna, 2010) or *C. elegans* (Gatfield *et al*, 2003). Most of mRNAs physiologically regulated by NMD belong to the long 3’-UTR category (Yepiskoposyan *et al*, 2011) but many mRNAs generated by point mutation in neoplastic cells belong to the EJC class (Lindeboom *et al*, 2016). Degradation of both types of NMD substrates requires three universally conserved “core” factors: Upf1, Upf2 and Upf3 (Leeds *et al*, 1992, 1991; Cui *et al*, 1995; Pulak & Anderson, 1993; Perlick *et al*, 1996). Independent of the initial triggering feature, RNA degradation in NMD occurs through activation of general RNA degradation pathways: decapping and endonucleolytic cleavage. Final degradation of RNA depends on the major 5’ to 3’ exonuclease Xrn1 (He & Jacobson, 2001) and the cytoplasmic exosome 3’ to 5’ exonucleolytic activity (Eberle *et al*, 2009).

How the NMD machinery differentiates normal termination codons from PTC has been a long-standing question (He & Jacobson, 2015). For long 3’-UTR NMD substrates, the prevalent model proposes that detection of a stop codon as premature occurs when the stop is far from the poly(A) tail, it is the faux 3’-UTR model(Amrani *et al*, 2004). In this model, the polyA-binding protein, Pab1 in yeast and PABPC1 in mammals, affects translation termination efficiency. Slow termination allows recruitment of Upf1, Upf2, Upf3, which leads to rapid decapping and degradation of NMD substrates (Muhlrad & Parker, 1994, 1999). A long distance from the PTC to PABPC1 triggers NMD in all studied species (Bühler *et al*, 2006; Eberle *et al*, 2008; Singh *et al*, 2008; Huang *et al*, 2011).

EJC-enhanced NMD does not require a long 3’-UTR and depends on an exon-exon junction located at more than 50 nucleotides downstream the PTC (Holbrook *et al*, 2004; Maquat, 2004). EJC positioning is an important element of the prevalent model for the first steps in mammalian NMD, called SURF/DECID. This model involves the formation of an initial complex that contains a ribosome stalled at a termination codon, the translation termination factors eRF1 and eRF3, Upf1 and the protein kinase Smg1 (SURF). Recruitment of the Upf2 and Upf3 proteins to SURF *via* a proximal EJC, leads to formation of the DECID complex in which Upf1 N and C-terminal regions are phosphorylated by Smg1 (Kashima *et al*, 2006). Phosphorylated Upf1 binds the Smg6 endonuclease and the Smg5-7 heterodimer, which, in turn, activates RNA deadenylation and decapping. Recruitment of a phosphatase ensures the return of Upf1 to its initial state for another NMD cycle (Ohnishi *et al*, 2003).

While the SURF/DECID model offers an explanation for EJC-enhanced NMD, it does not apply to organisms that do not have EJC components, such as the budding yeast, and does not explain how NMD works on RNAs with long 3’- UTRs. Understanding how such substrates are degraded through NMD is relevant for mammalian cells since several physiologically important NMD substrates do not depend on an EJC downstream the termination codon. A salient example is the mRNA of the GADD45A gene (Chapin *et al*, 2014; Lykke-Andersen *et al*, 2014), a transcript whose destabilization through NMD is essential for the development of fly and mammal embryos (Nelson *et al*, 2016). It is possible that EJC-enhanced and EJC-independent NMD are two versions of the same, not yet understood, generic NMD mechanisms. Such a mechanism is unlikely to rely on a SURF/DECID-like sequence of events for several reasons. First, even if most of the RNA decay factors, and the key NMD proteins are conserved from yeast to humans, the SURF/DECID model proposes crucial roles for factors or events that are not conserved in all eukaryotes. For example, the protein kinase Smg1, a central component of the SURF complex, has no known equivalent in *S. cerevisiae* and its absence has little impact on NMD in *D. melanogaster* embryos (Chen *et al*, 2005). Second, a major assumption of the SURF/DECID model, the role of EJC in bringing Upf3 in the proximity of Upf1 (Le Hir *et al*, 2001; Chamieh *et al*, 2008), was challenged by very recent biochemical analyses that show interactions of Upf3 with translation termination complexes independent of EJC components (Neu-Yilik *et al*, 2017).

Third, while all the RNA degradation mechanisms can play a role in NMD, the model leaves open the question of the mechanisms of recruitment for the RNA degradation factors. Major roles have been attributed to the decapping machinery (Muhlrad & Parker, 1994; Lykke-Andersen, 2002), endonucleolytic cleavage (Huntzinger *et al*, 2008; Eberle *et al*, 2009) or deadenylation (Loh *et al*, 2013), depending on the reporter RNA used. The molecular mechanisms responsible for the redundancy and interplay between these different degradation pathways (Metze *et al*, 2013; Colombo *et al*, 2016) remain unclear. Hence, a re-evaluation of the available data and new biochemical results on the involved complexes are critical to uncover a universally conserved mechanism of NMD in eukaryotes. Such a mechanism could then serve as the basis to understand the evolution of NMD, its variations across species and additional levels of complexity.

The enumerated weaknesses of current NMD models originate, at least in part, from the lack of a global view of the biochemistry of the NMD process. We show here that fast affinity purification of yeast NMD complexes coupled with high-resolution quantitative mass spectrometry (LC-MS/MS) allows a global and unprecedented view of NMD complexes. Our results uncover the existence of two distinct and successive protein complexes that both contain the essential NMD factor Upf1. We identified two yeast accessory NMD factors that interact with Upf1 and become essential for NMD under limiting conditions. Their similarity with Smg6 and Smg5-7 implies that comparable molecular mechanisms for NMD have been conserved from yeast to humans. Furthermore, our results suggest that Upf1 might suffer major conformation changes to accommodate a switch from a “Detector” complex to an “Effector” complex, that triggers RNA decapping, the first step in NMD substrates degradation.

## Results

### Quantitative mass spectrometry reveals the global composition of Upf1 associated complexes

Previous large-scale attempts to describe the composition of Upf1 associated complexes in yeast failed to identify known Upf1 partners, such as Upf2 or Upf3 (Gavin *et al*, 2006; Krogan *et al*, 2006; Collins *et al*, 2007) even if these studies provided high-quality interaction data on other proteins. This negative result could be due to the relatively low abundance of Upf2 and Upf3, estimated to 1470 and 1439 molecules per cell (Ho *et al*, 2018), which places them in the category of the 25% least abundant proteins in yeast (**Fig. 1a**). The instability of the complexes during the purification procedure could also play a role. To overcome these problems we used a combination of fast affinity purification (Oeffinger *et al*, 2007) with the current high sensitivity of mass-spectrometry methods for protein identification. TAP-tagged (Rigaut *et al*, 1999) Upf1 was isolated from cells extracts obtained from 6L of yeast culture grown in exponential phase. The total cell extract was used as a reference and an untagged strain as a negative control. Four hundred proteins were reliably identified in the total extract and corresponded, as expected, to abundant yeast proteins (the frequency distribution of the abundance for the identified proteins is shown as dark grey bars in **Fig. 1b**). Many abundant proteins were also identified in the Upf1-TAP purified fraction (distribution in dark grey, **Fig. 1c**). In contrast to the input fraction, less abundant proteins were also identified in the purified fraction (distribution in light grey, **Fig. 1c**).

**Fig. 1:**
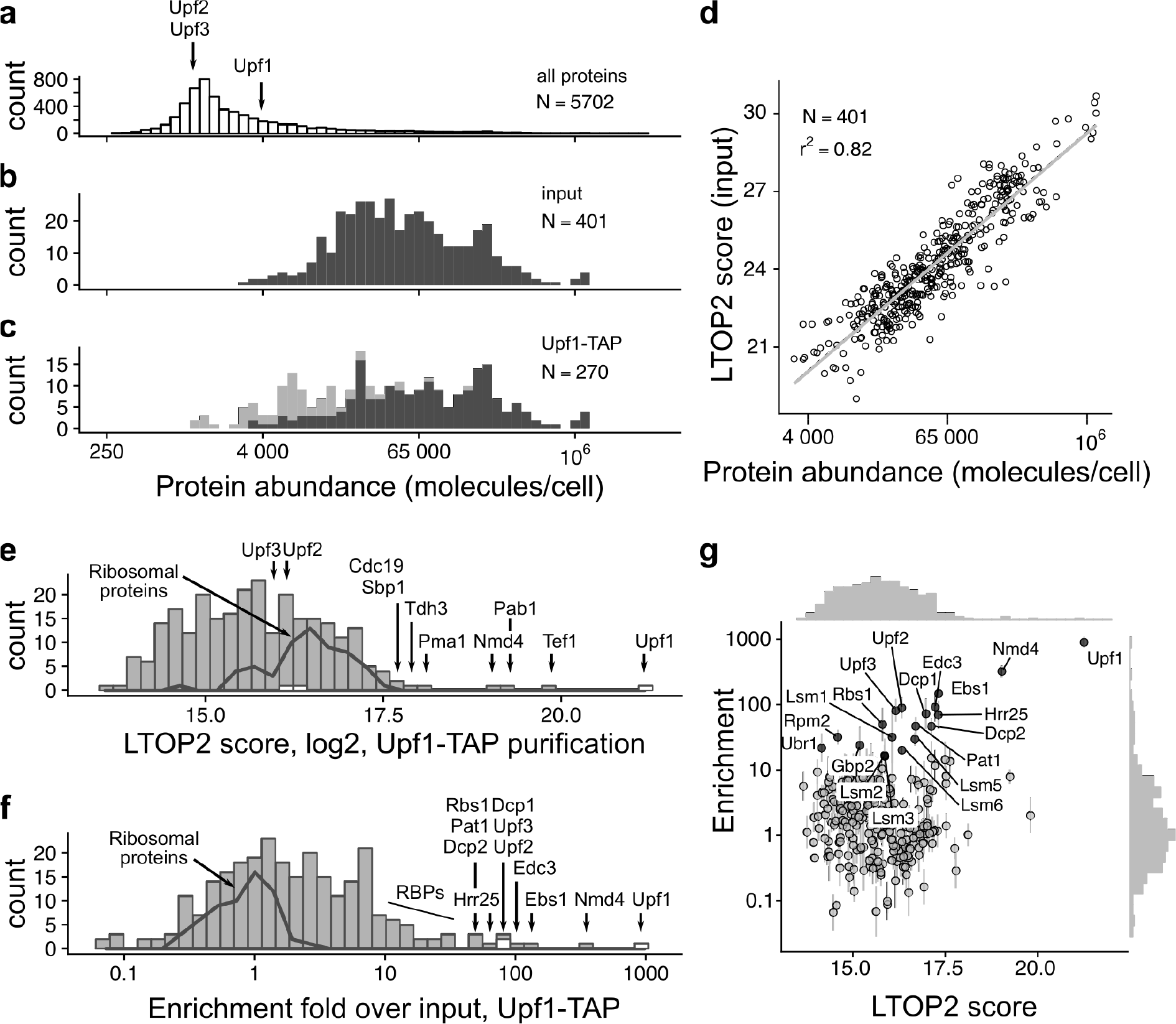
Enrichment analysis accurately describes Upf1-associated proteins. **a.** Positioning of Upf1, Upf2 and Upf3 in the abundance distribution of all yeast proteins, based on the data compiled by Ho et al., 2018. The distribution of abundance for the proteins identified by mass spectrometry in input (**b**), and purified samples (**c**), show the expected bias for proteins present in high copy numbers. Light grey colour highlights the positions of proteins that were only identified in the Upf1-TAP samples (6 replicates) and not in the total extract (3 replicates). **d.** Correlation of LTOP2-based protein level estimates in a total yeast protein extract with published protein abundance data (Ho et al., 2018). **e.** Distribution of the intensity for the proteins quantified in association with Upf1 (LTOP2 score, log scale). Positioning of Upf1, Upf2 and Upf3 are highlighted as white rectangles. The dark grey line indicates the region of intensities corresponding to ribosomal proteins. **f.** Distribution of the protein enrichment values for Upf1 associated proteins based on LTOP2 scores and known protein abundance in a total yeast extract. Upf1, Upf2 and Upf3 positions are also highlighted as white rectangles. **g.** A combination of the data presented in (**e**) and (**f**) as a scatter plot, to see both the amount and enrichment of proteins in Upf1-TAP purified samples. The horizontal axis represents the LTOP2 score, with the vertical axis showing enrichment over input values. Both axes use log transformed values.

To assess to what extent the identified proteins were specific to the Upf1 purification, we used a label-free quantification method based on the intensity of the most abundant peptides for each of the identified proteins (Silva *et al*, 2006), a method that accurately estimates protein levels (Ahrné *et al*, 2013). For each identified protein, we derived a peptide intensity-based score that we called LTOP2 - “L”, because it includes a log2 transformation, and “TOP2” for the minimum number of required peptides (see Methods and Supplementary Table 1). The LTOP2 score of proteins identified in the total protein extract was well correlated with protein abundance estimates (Ho *et al*, 2018), with a Pearson determination coefficient of 0.82 (**Fig. 1d**).

Upf1 had the highest LTOP2 score in the Upf1-TAP purification, as expected, validating both the use of this MS score for protein quantitation and the efficiency of the affinity purification. Most of the other proteins with high LTOP2 scores in the purification, such as the plasma membrane proton pump Pma1, or the glyceraldehyde-3-phosphate dehydrogenase Tdh3 (**Fig. 1e**), were unrelated to Upf1 or NMD, but are among the 10% most abundant proteins in yeast. NMD factors Upf2 and Upf3, the best known partners of Upf1, ranked low in the list of 270 LTOP2 scores, with position 80 and 91 respectively (Supplementary Tables 1 and 2). Thus, many of the proteins present in the affinity purified sample were contaminants, known to pollute affinity purifications and complicate the conclusions of such experiments (Mellacheruvu *et al*, 2013). To filter this type of contamination, we chose to calculate an enrichment score, which reflects the ratio between the proportion of a protein in a sample and its proportion in a total extract (Ho *et al*, 2018). Top enrichment scores corresponded to the tagged protein Upf1, Upf2 (rank 5) and Upf3 (rank 6), validating enrichment as an excellent noise removal strategy in the identification of specific partners in affinity-purified samples (**Fig. 1f**, and **Supplementary Tables 3 and 4**). To simultaneously visualize both the amount of protein found in a purified sample and its enrichment score, we combined LTOP2 and enrichment in a graphical representation that is similar to the intensity versus fold change “MA” plots used for the analysis of differential gene expression (**Fig. 1g**).

In addition to the canonical Upf1-Upf2-Upf3 interactions (He *et al*, 1996, 1997), our enrichment strategy identified the RNA decapping enzyme Dcp2, its cofactor Dcp1, and decapping activators, Edc3, Lsm1-7, and Pat1 as Upf1 partners. This result was correlated with the requirement for decapping for the degradation of NMD substrates in yeast (Muhlrad & Parker, 1994). Direct or indirect interactions of Upf1 with these different factors have been previously reported, mostly through yeast two-hybrid, or affinity purification coupled with immunoblots (He & Jacobson, 2015). Unexpectedly, two poorly characterized proteins, Nmd4 and Ebs1 ranked second and third in the Upf1 purification. Both proteins were previously linked with NMD. Nmd4 was identified in yeast Upf1 two-hybrid screen but was not further studied(He & Jacobson, 1995). Ebs1, identified through sequence similarity as a potential yeast equivalent of human Smg7, a component of mammalian NMD, was shown to co-purify with Upf1 and to affect yeast NMD (Luke *et al*, 2007). Other proteins enriched in the Upf1-associated fraction by a factor of 10 or more (Rbs1, Hrr25, Gbp2, Gar1, Cbf5, Sbp1, Ssd1, Lsm12, Lhp1 and Rbg1) have functions related with RNA metabolism and could be indirectly associated with the Upf1 RNA-containing complexes. None of the proteins found enriched with Upf1 by a factor of 8 or higher were detected in the control purification (Supplementary Tables 3 and 4). These results indicated that Upf1 has more protein partners than previously thought and validated quantitative mass-spectrometry of purified complexes as a powerful strategy to characterize yeast NMD complexes.

### Upf1-associated proteins are grouped in two mutually exclusive complexes

Since our data indicated that Upf1 is associated with a relatively large number of factors, and since Upf1 directly binds RNA (Weng *et al*, 1998) in large polysomal complexes (Atkin *et al*, 1997), some identified interactions could be mediated by RNA. To establish which interactions were RNA dependent, we repeated the Upf1-TAP purification, in a protocol including two RNase A and RNase T1 treatments. Ribosome constituents and RNA binding proteins were lost following this harsh RNase treatment (p-values associated with GO term enrichment analysis between 10^−7^ and 10^−30^ for the corresponding terms, **Supplementary Table 5**). RNase treatment was efficient, suggesting that the proteins that remained specifically enriched were bound to the Upf1 complex through direct protein-protein interactions. Among these proteins, we identified the decapping enzyme Dcp2 and its co-factors Dcp1 and Edc3, Nmd4, Ebs1, and the protein kinase Hrr25, a homologue of mammalian casein kinase delta, CSNK1D (**Fig. 2a**). Upf2 and Upf3 interactions were sensitive to RNase. In view of these results, and to obtain a global view of NMD complexes in yeast, we affinity purified TAP-tagged versions of the components of Upf1-associated complexes identified here with or without RNase treatment: Upf2, Upf3, Nmd4, Ebs1, Dcp1, and Hrr25 (**Supplementary Table 4**).

**Fig. 2:**
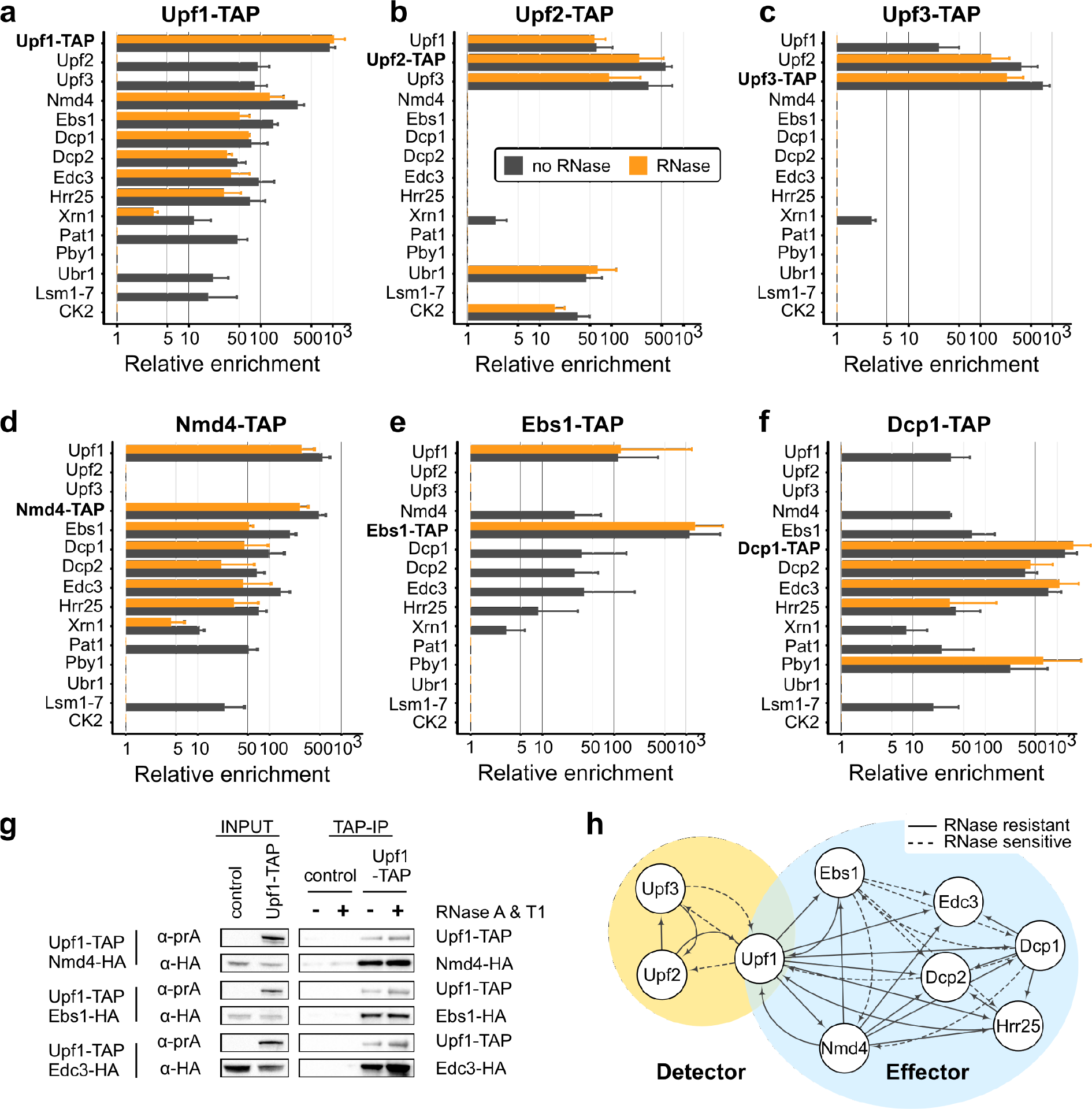
Purification of NMD factors reveals two distinct complexes containing Upf1. Enrichment values for proteins identified in tagged Upf1 (**a**), Upf2 (**b**), Upf3 (**c**), Nmd4 (**d**), Ebs1 (**e**), and Dcp1 (**f**) purifications. Tagged proteins are indicated in bold for each experiment. Black bars correspond to enrichment values obtained in purifications done without RNase and orange bars with an RNase A and RNase T1 treatment. Only proteins enriched by a factor of 16 or more in one of the purifications presented here are shown. For clarity, groups of related proteins were combined (the group of Lsm1 to Lsm7 is marked as “Lsm1-7” and the values correspond to the mean of all the values found in purifications; CK2 group corresponds to Cka1, Cka2, Ckb1 and Ckb2). **g.** Immunoblot validation of Upf1-TAP interactions with HA tagged Nmd4, Ebs1 and Edc3. Control strains did not express Upf1-TAP. **h.** Representation of binary interactions identified by our experiments and which define two complexes containing Upf1: Detector and Effector; dashed lines correspond to interactions that are sensitive to RNase treatment; the arrows start at the tagged protein and indicate the enriched factor.

One of the most surprising results of the extensive purification data was the difference between the sets of proteins enriched with Upf2-TAP (**Fig. 2b**) or Upf3-TAP (**Fig. 2c**) in comparison with Upf1-TAP (**Fig. 1g** and **2a**). Many of the proteins present in the Upf1 complex, such as the decapping factors, Ebs1, and Nmd4 were absent in the Upf2-TAP and Upf3-TAP purifications. Conversely, the purifications done with Nmd4-TAP, Ebs1-TAP, Dcp1-TAP or Hrr25-TAP, did not co-purify Upf2 or Upf3, even if Upf1 was specifically enriched with or without RNase treatment (**Fig. 2d, e, f** and **Supplementary Table 4**). Nmd4-TAP shared the same purification profile with Upf1-TAP except for Upf2 and Upf3 (**Fig. 2a, d**). Ebs1-TAP, also purified the same set of proteins, but the enrichment of most of the factors, except Upf1, was dependent on RNA (**Fig. 2e**). Dcp1 co-purified with its known partners Dcp2 and Edc3, but also with Upf1, Nmd4, Ebs1, and Hrr25 (**Fig. 2f**). In addition to Upf1-associated proteins, purifications with Dcp1-TAP and Hrr25-TAP revealed their presence in other protein complexes. For example, Dcp1 purification also showed the enrichment of another decapping-associated factor, Pby1 (**Fig. 2f**). The results, including the variability of biological replicates, are presented in **Supplementary Tables 3** and **4** and can be explored at hub05.hosting.pasteur.fr/NMD_complexes. To confirm the mass-spectrometry quantitative results by a different detection method, we purified Upf1-TAP in strains in which Nmd4, Ebs1 and Edc3 were tagged with 3 repeats of the HA epitope at the C-termini. Immunoblots on purified fractions showed that these three proteins were enriched with Upf1 with or without RNase treatment, as expected (**Fig. 2g**).

The obtained results suggest that Upf1 is part of two mutually exclusive complexes, one composed of Upf1, Upf2 and Upf3 and the other containing Upf1, Nmd4, Ebs1 and decapping factors (**Fig. 2h**). Since the second complex contains the decapping machinery, known to trigger the degradation of NMD substrates (Muhlrad & Parker, 1994), we refer to this complex as the “**Effector**”. It is likely that the action of Upf2 and Upf3 precedes decapping and ensures specificity for NMD substrates, so we decided to call the Upf1-Upf2-Upf3 complex, the “**Detector**”. Altogether, these data suggest that Upf1 coordinates the recognition and degradation of NMD by binding to two mutually exclusive sets of proteins.

### The Upf1 cysteine-histidine rich N-terminal domain is an interaction « hub » *in vivo*

To find out how the NMD complexes are organised and to investigate why the Upf1-binding proteins in Detector and Effector were mutually exclusive, we tested the importance of Upf1 domains in the interaction with the associated factors. We built plasmids for the expression of tagged Upf1 fragments (**Fig. 3a**) to test the cysteine-histidine rich N-terminal domain (Upf1-CH, 2-208), the C-terminal helicase domain (Upf1-HD-Cter, 208-971), as well as variants of these constructs, comprising either the helicase domain alone (Upf1-HD, 208-853) or a truncated Upf1 lacking the C-terminal extension (Upf1-CH-HD, 2-853). Strains deleted for endogenous *UPF1* were transformed with plasmids expressing N-terminal TAP-tagged full length Upf1 (Upf1-FL) or the various fragments. TAP-Upf1-FL was expressed to levels ten to fifty folds higher than the C-terminal tagged Upf1 expressed from the chromosomal locus, as estimated by immunoblots on total protein cell extracts. The various fragments were also stably expressed to high levels at the expected size (**Supplementary Fig. 2a**).

**Fig. 3:**
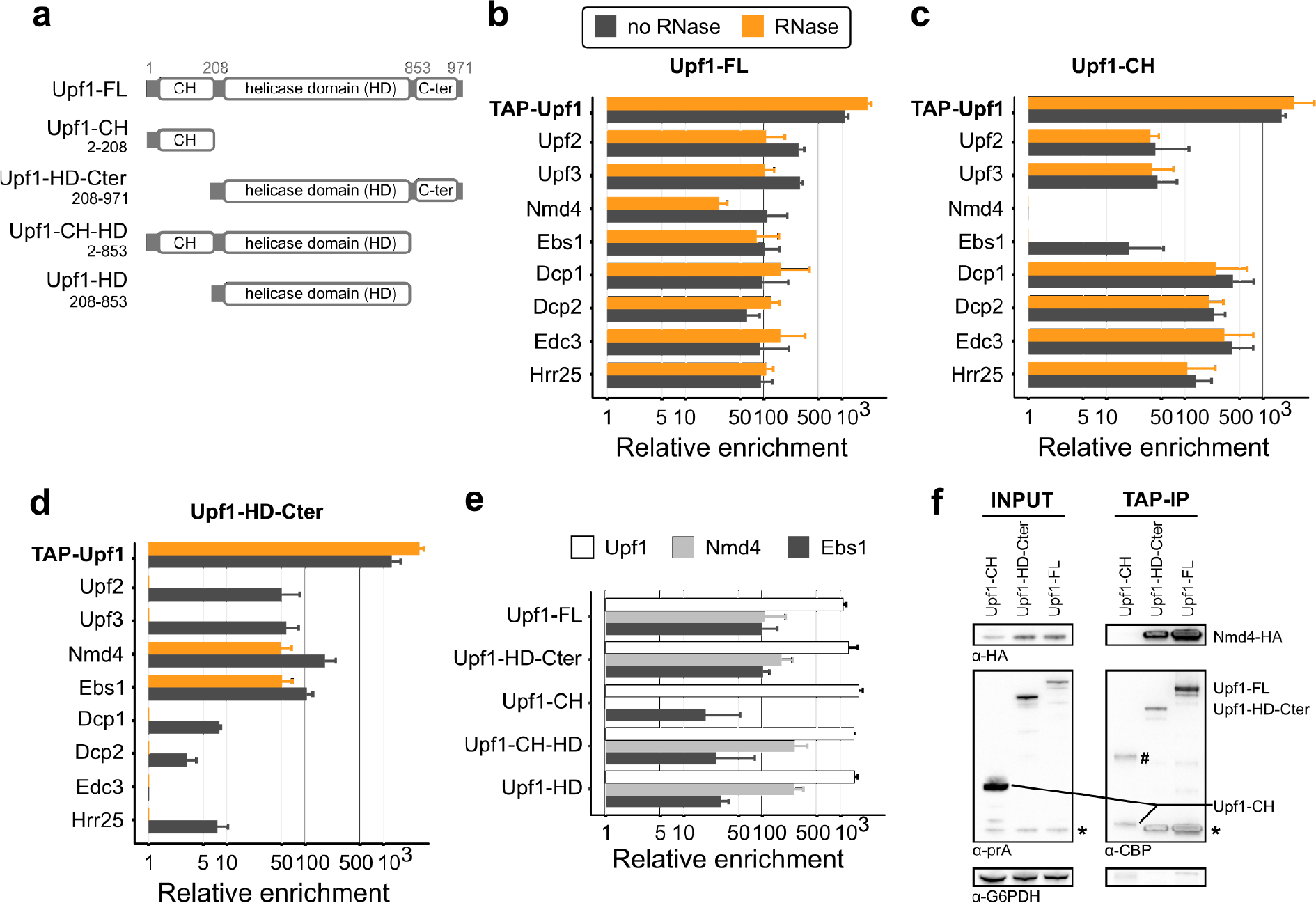
Nmd4 and Ebs1 are the only Detector components that interact with Upf1 independent of the N-terminal CH domain. **a.** Upf1 fragments tested: FL is for the full-length protein; HD-Cter for the region containing the helicase domain and the C-terminal part of Upf1 (208-971); CH for the N-terminal domain (2-208) that contains the N-terminal unstructured region and the Cysteine-Histidine rich domain; CH-HD for a version a full-length Upf1 lacking the C-terminal region 854-971; and HD for the helicase domain of Upf1 alone (not to scale). N-terminal TAP-tagged Upf1 versions were expressed from a single copy vector under an inducible tetO7 promoter. **b.** to **d.** Results of purifications using Upf1-FL (**b**), Upf1-CH (**c**) and Upf1-Cter (**d**) as bait with the x-axis showing the enrichment value (log scale) for Effector and Detector components. The colour of the bars illustrates the treatment used during the purification, black without RNase and orange with an RNase treatment. **e.** Comparison of the enrichment of Upf1, Nmd4 and Ebs1 in the purification of the Upf1 fragments. The Upf1 C-terminal region (854-971) affected the association with Nmd4 or Ebs1 (Student t-test with a one-sided alternative hypothesis. We compared the enrichment of each of Ebs1 and Nmd4 between purifications of Upf1 fragments having this region (Upf1-FL and Upf1-HD-Cter, 6 experiments) and purifications of Upf1 fragments lacking the extension (Upf1-CH-HD and Upf1-HD, 4 experiments). The C-terminal region had no effect on Nmd4 enrichment (p-value ≈ 0.98), whereas there was significantly less associated Ebs1 on this small region (p-value ≈ 0.0013). **f.** Western blot of input and eluates of Upf1 domains purification in a Nmd4-HA strain. The band with the # might corresponds to a dimer of Upf1-CH, bands marked with a star correspond to residual signal with the anti-HA antibodies (Nmd4). Fragments in the eluate have a smaller size because the protein A part of the tag was removed by digestion with the TEV protease. G6PDH served as a loading control in the input samples.

Enrichment of the various proteins in the purification done with overexpressed TAP-Upf1-FL was well correlated with the enrichment of the same proteins in the previous purifications done with the protein tagged at its C-terminus and expressed from the endogenous *UPF1* locus (**Supplementary Fig. 2b and Supplementary Table 4**), confirming that neither overexpression nor tag position had major effects on Detector and Effector composition (see **Fig. 2a** compared with **Fig. 3b**). The only marked difference between the results with the overexpressed N-terminal Upf1 and the C-terminal tagged Upf1 was the persistence of Upf2 and Upf3 among the enriched proteins after RNase treatment (**Fig. 3b**, orange bars). Since the Upf1 interaction was also preserved when the Detector complex was purified using tagged Upf2 (**Fig. 2b**), but not when using tagged Upf3 (**Fig. 2c**), this variability in Detector sensitivity to RNase indicates the importance of RNA for the stability of the Upf1-Upf2-Upf3 complex. In contrast, the interactions of Upf1 with the Effector complex proteins, and especially with Nmd4, were insensitive to RNase in all the studied situations.

Tagged Upf1-CH, recovered a majority of the proteins associated with the full-length protein, with the notable exception of Nmd4 (not detected) and of Ebs1, which was detected only in the sample not treated with RNase, albeit to a lower intensity than in the full-length Upf1 purification (**Fig. 3b, c**). The observation that the other proteins of the Effector and Detector complexes were highly enriched in this purification and the fact that the interactions were not sensitive to RNase (**Fig. 3c**), raise the possibility that Upf2 and Upf3 compete with the decapping factors for binding to Upf1-CH domain. Thus, the CH domain seems to be crucial for both Detector and Effector organisation.

### Nmd4 and Ebs1 are tightly associated with Upf1-HD-Cter

Nmd4 and Ebs1 were the only proteins interacting specifically with the HD-Cter domain of Upf1 independently of RNA (**Fig. 3d**). To establish if the C-ter domain of Upf1 could play a role in these interactions, we purified Upf1-CH-HD and Upf1-HD and evaluated the relative enrichment levels for Ebs1 and Nmd4 in these conditions. While Nmd4 was recovered with both Upf1 fragments, Ebs1 levels were substantially lower in the absence of the C-terminal extension (**Fig. 3e**, t-test p-value < 0.001). These results suggest that the C-terminal extension of Upf1 has a crucial effect on binding Ebs1, and does not affect binding of Nmd4.

To confirm the observed strong Nmd4-Upf1 interactions, we built Nmd4-HA strains and tested the presence of the tagged protein in fractions enriched with overexpressed tagged Upf1-FL, Upf1-CH and Upf1-HD-Cter. The purification of Upf1-FL and Upf1-HD-Cter co-enriched Nmd4-HA, while the Upf1-CH region alone did not. Unexpectedly, overexpression of Upf1-FL or Upf1-HD-Cter also led to increased levels of Nmd4-HA in the total extracts (**Fig. 3f**), suggesting that Nmd4 protein stability can be affected by its interaction with Upf1. To test the importance of the observed interaction between Nmd4 and Upf1 for the association with other factors of the Effector complex we purified Upf1-TAP in the absence of *NMD4.* We observed no major effects on the various associated proteins (**Fig. 4a**, **Supplementary Fig. 3** and **Supplementary Table 4**). In the reciprocal experiment, in which we purified the Nmd4-associated complexes in the absence of *UPF1*, all the specific components of the Effector complex were lost (**Fig. 4b**). These results, together with the strong and RNase insensitive enrichment of Nmd4 in Upf1 complexes, suggested that Upf1 directly interacts with Nmd4. To validate this hypothesis, we expressed CBP-Nmd4 as a recombinant protein in *E. coli* and tested its association *in vitro* with recombinant purified Upf1 helicase domains of yeast and human origin (yUpf1-HD 220-851 and hUpf1-HD 295-914, **Fig. 4c**). Yeast Upf1, but not human Upf1 co-purified with CBP-Nmd4, showing that the interaction is direct and specific (**Fig. 4d**). These results designate Nmd4 as the most tightly bound Upf1 co-factor, specific to the conformation of Upf1 present in the Effector complex.

**Fig. 4:**
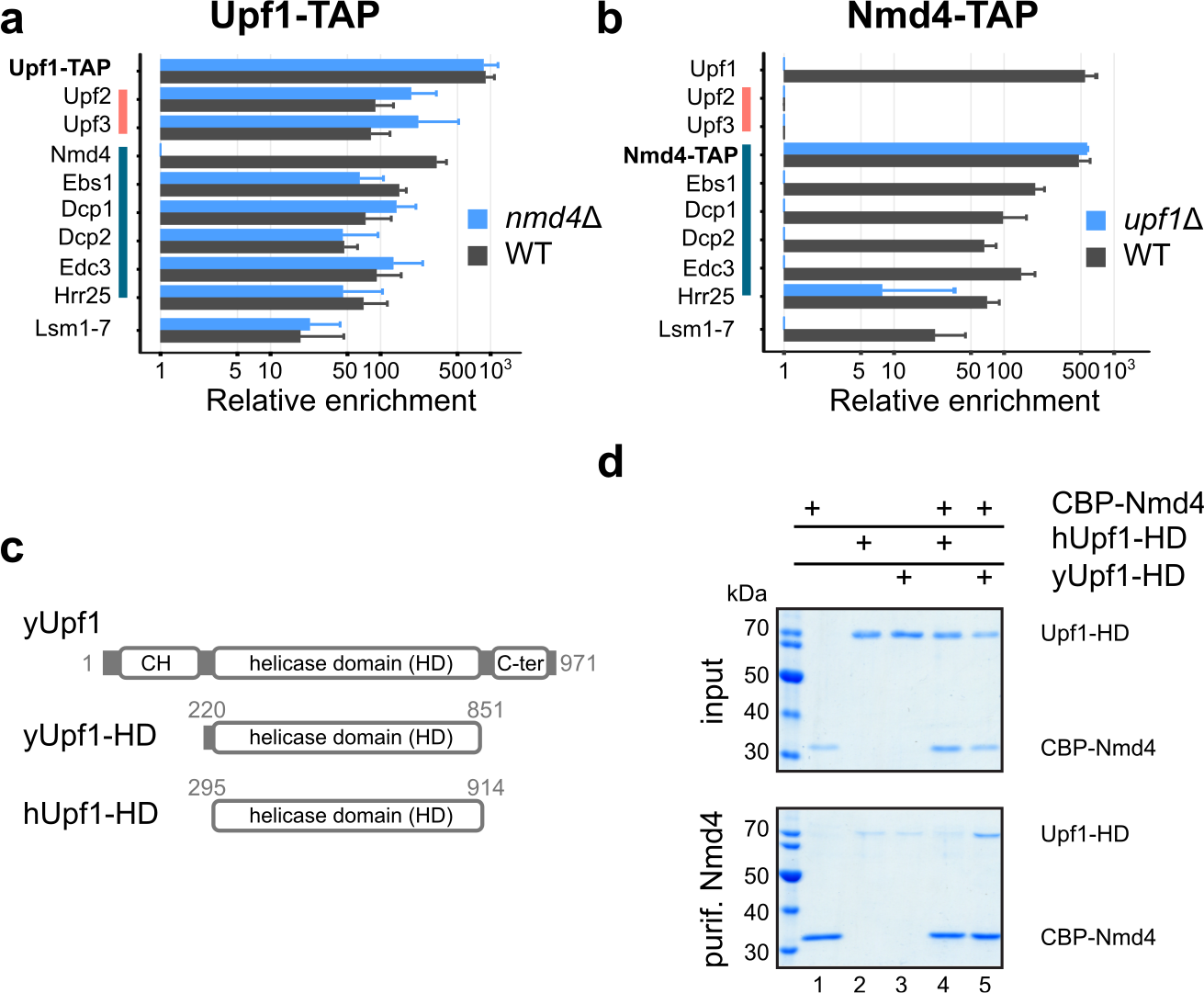
Nmd4 interaction with the Effector complex is direct and mediated by Upf1. **A.** Enrichment of Detector and Effector components in purified Upf1-TAP in the presence (black bars) and absence (blue bars) of NMD4. Pink and dark blue vertical lines highlight proteins of the Detector and Effector complexes respectively. **B.** Enrichment of Effector components in Nmd4-TAP in the presence (black bars) and absence (blue bars) of UPF1. **C.** Schematics of Upf1 fragments used for the *in vitro* interaction assay; yUpf1-HD is the helicase domain (220-851) of the yeast Upf1 protein, hUpf1-HD is the helicase domain (295-914) of human Upf1 (not to scale). **D.** CBP-Nmd4 was mixed with hUPF1-HD or yUPF1-HD (all the proteins overexpressed in *E. coli* and purified). Protein mixtures before (input, 20% of total) or after purification on calmodulin affinity beads were separated on 10% SDS-PAGE (w/v) acrylamide gels.

### Potential Smg5-6-7 homologs can be essential for NMD efficiency under limiting conditions

The association of Nmd4 and Ebs1 to Upf1 support the hypothesis that they are the yeast functional equivalents of human Smg6 and Smg5-7. In line with this idea, Nmd4 contains an endonuclease-like region from the PIN domain family (Clissold & Ponting, 2000), like Smg6 and Smg5 (**Fig. 5a**). Ebs1 has strong similarities with the N-terminal 14-3-3 domains of Smg5, Smg6 and Smg7 (Luke *et al*, 2007), with similar percentages of identity in the aligned sequences for this domain (**Fig. 5a**, **Supplementary Fig. 4**). The interactions that we described here indicate that Nmd4 and Ebs1 are components of the Effector complex and co-factors of Upf1. While a role for Ebs1 in the degradation of an NMD reporter RNA has been previously reported (Luke *et al*, 2007), no data were available about a potential global role of *NMD4* and *EBS1* in NMD. To investigate the impact of *NMD4* and *EBS1* on NMD on a large scale, we performed strand-specific RNASeq experiments in strains deleted for each of the two genes.

**Fig. 5:**
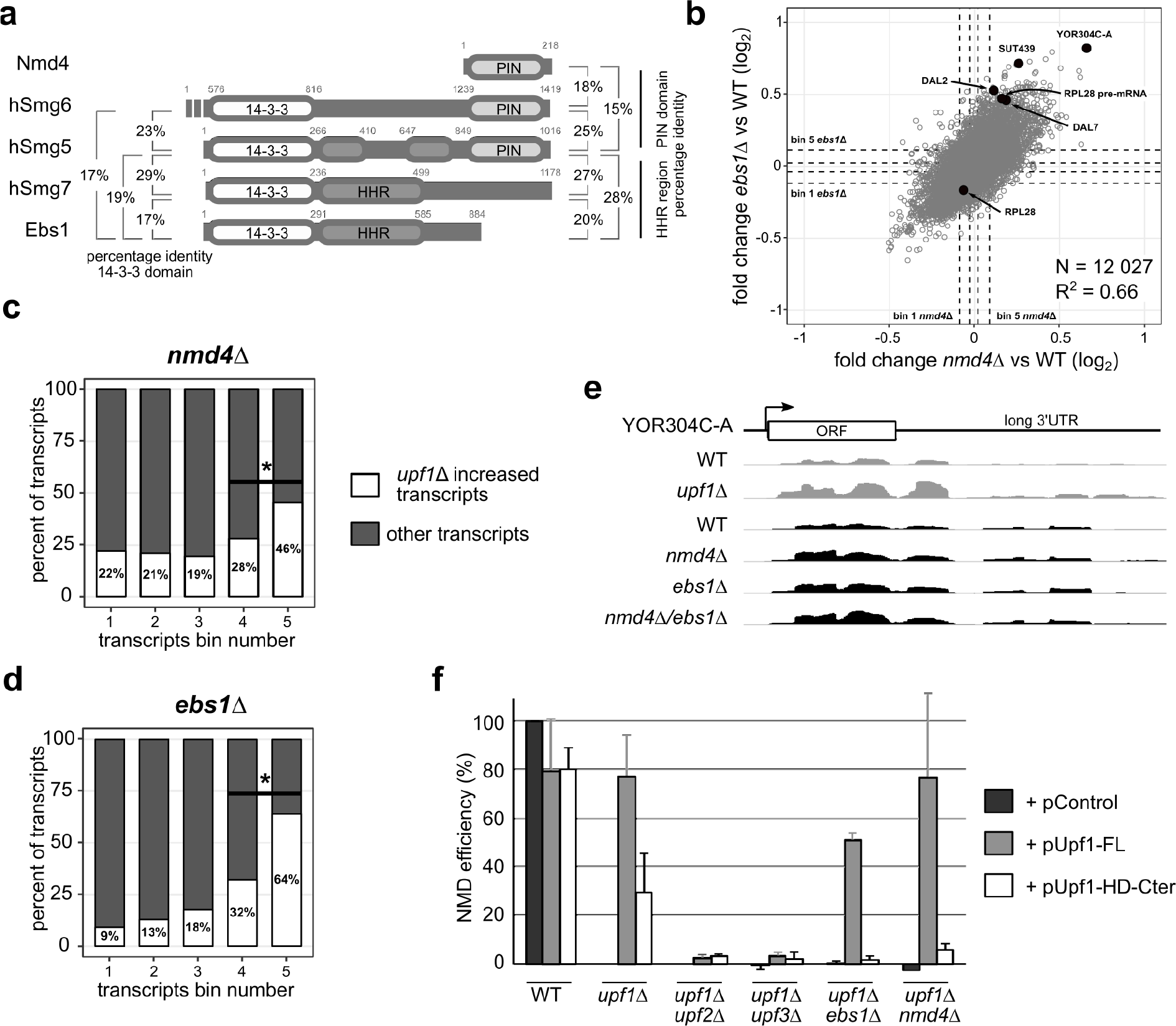
Nmd4 and Ebs1 are essential for NMD elicited by the overexpression of the helicase domain of Upf1. **a.** Schematic representation of the domain structure of Nmd4 and Ebs1 from *S. cerevisiae* compared with human (h) Smg6, Smg5 and Smg7. 14-3-3, HHR and PIN domains were defined based on literature data (Fukuhara et al., 2005, Luke et al., 2007). Boundaries of the PIN domains were chosen based on a Mafft alignment of Smg5-6 of several species, while for Nmd4 the entire protein sequence was used. Drawing is not to scale. Identity percentages among the different domains of Smg proteins, Nmd4 and Ebs1 are indicated. Values represent percent of identical residues as a fraction of all the aligned residues. **b.** Scatter plot of the mean fold change of transcripts in *nmd4*Δ RNAseq experiment against transcript mean fold change in *ebs1*Δ (Pearson correlation, r = 0.66, p-value < 2.10^−16^). Mean fold changes were computed using DESeq2 with BY4741 as reference, with 3 replicates for each condition. Black dots correspond to NMD substrate examples for which individual traces are shown in Figure S4. For RPL28, we highlighted the pre-mRNA, NMD sensitive, and the mature form that is not. Dashed lines represent boundaries of five bins of equal numbers of transcripts (2046 transcripts or notable features per bin, see the STAR Methods), with bin 5 containing the transcripts that were most increased in the mutant condition compared with wild type. **c.** Percentage of transcripts affected by *UPF1* deletion (increase by at least 1.4 fold) among *nmd4*Δ bins, as defined in **b**. Differences between percentages of *upf1*Δ affected transcripts in bin 4 and 5 and bin 3 and 4 were significant (binomial test, p<10^−24^). **d.** Same as in **c**, for *ebs1*Δ. Differences between distribution of transcripts in bin n and bin n+1 were significant (binomial test, p<10^−9^). **e.** Example of YOR304C-A up regulated in *nmd4*Δ, *ebs1*Δ and the double mutant *nmd4*Δ/*ebs1*Δ. **f.** NMD efficiency of WT, *upf1*Δ or double mutant strains complemented with Upf1-FL, Upf1-HD-Cter or an empty plasmid. The NMD efficiency for each strain is based on reverse-transcription followed by quantitative PCR for RPL28 pre-mRNA, an abundant NMD substrate. A wild-type strain has 100% NMD efficiency and a *upf1*Δ strain has 0% NMD efficiency, by definition.

The obtained RNASeq data allowed us to quantify variations of RNA levels for 12 027 transcripts (mapping statistics available in **Supplementary Table 6**). DESeq2 (Love *et al*, 2014) was used to adjust, normalize and compute the changes in expression levels in *ebs1*Δ and *nmd4*Δ strains in comparison with a wild-type strain. Even if the amplitude of change in transcript levels was modest, the observed variations were highly correlated between the two strains (**Fig. 5b**, N = 12 027; Pearson correlation coefficient = 0.66; p-value < 2.2 10^−16^). Examples of observed transcript changes are presented in **Supplementary Fig. 5**. To see to what extent the transcripts enriched in these mutant strains were related with transcripts stabilized in the absence of *UPF1*, we performed a similar RNAseq experiment by comparing *upf1*Δ to a wild-type strain. 3 271 transcripts showed an increase of more than 1.4 fold in the absence of *UPF1* and were used as a reference set for NMD-sensitive RNAs (**Supplementary Fig. 5**). To estimate the presence of such RNAs in the population of transcripts affected by the absence of *NMD4* or *EBS1*, we divided the set of obtained values in five bins of equal size (N = 2 406 per bin, **Fig. 5b**). We found a strong enrichment of *UPF1*-affected RNA in bin 5, which contains the RNAs most increased in the absence of *NMD4* **(Fig. 5c**) or *EBS1* (**Fig. 5d**) (p-value < 10^−74^, binomial test performed with one-sided alternative null hypothesis, N = 2 406). Deletion of both *NMD4* and *EBS1* did not have a synergistic effect (see **Fig. 5e** and **S5** for examples of RNASeq results). As expected from the presence of both proteins in the Effector complex, this result suggests that *NMD4* and *EBS1* affect similar, and not parallel, steps in NMD.

To establish how *NMD4* and *EBS1* could affect NMD substrates, we searched for conditions that would simplify our phenotypic analysis. We chose to focus on the helicase and C-terminal domain of Upf1, important for binding of both Nmd4 and Ebs1 (**Fig. 3d**). It has been previously observed that overexpression of this region of Upf1 can complement the deletion of the *UPF1* gene (Weng *et al*, 1996). We thus tested whether *NMD4* and *EBS1* were required for this complementation. First, we validated the effect of overexpressing full-length Upf1 or its different domains (depicted in **Fig. 3a**) on the levels of RPL28 pre-mRNA, a well-studied NMD substrate. To facilitate the visualization of the complementation levels, we devised an NMD efficiency measure that takes into account RPL28 pre-mRNA levels in WT (100% NMD) and *upf1* (0% NMD) strains (see **Methods**). While overexpression of full-length *UPF1Δ* led to the destabilization of RPL28 pre-mRNA, with an estimated reconstituted NMD efficiency of 80%, overexpression of Upf1-HD-Cter allowed recovery of 30% of NMD in a *upf1Δ* strain (**Fig. 5f** and **Supplementary Fig. 6**). Importantly, deletion of either *EBS1* or *NMD4* abolished the partial complementation effect (**Fig. 5f**), suggesting that under limiting conditions, *EBS1* and *NMD4* become essential for NMD. These results correlated with the inability of the Upf1-HD fragment, which showed decreased binding of Ebs1 (**Fig. 3e**), to complement NMD (**Supplementary Fig. 6**). In line with these observations, *EBS1* deletion also decreased the NMD efficiency of Upf1-FL (**Fig. 5f**). Altogether, these results indicate that the mechanism by which Upf1-HD-Cter destabilizes NMD substrates depends on the presence of Nmd4 and Ebs1, the only factors that specifically interact with this region of Upf1 independent on RNA (**Fig. 3d**). To investigate if this effect was due to a canonical NMD mechanism, we performed the complementation assays for Upf1-HD-Cter in strains lacking *UPF2* or *UPF3*. No complementation could be observed in these conditions (**Fig. 5f**). These data suggest that Ebs1 and Nmd4 are specific partners of Upf1 helicase and C-terminal domain and play a crucial role in the function of this Upf1 region in NMD.

**Fig. 6:**
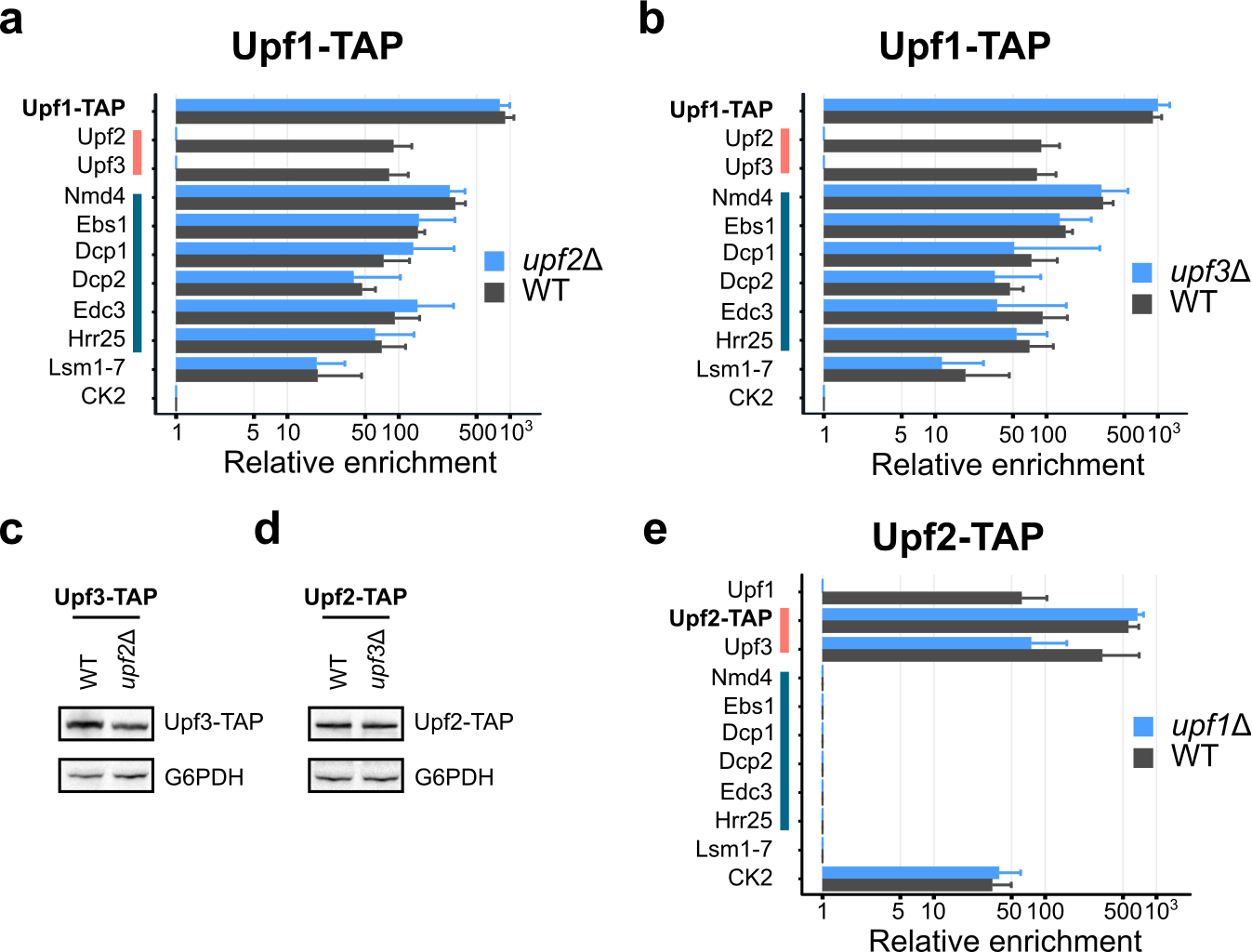
Upf2 and Upf3 function as a heterodimer in the formation of Detector. Comparison between enrichment values for Effector and Detector components, Lsm proteins and CK subunits in Upf1-TAP purified samples in the presence (grey bars) or absence (blue bars) of *UPF2* (**a**) or *UPF3* (**b**). Evaluation of the effects of *UPF2* deletion on Upf3-TAP levels (**c**) and of *UPF3* deletion on Upf2-TAP levels (**d**) in total extracts, in comparison with a loading control (G6PDH) was done by immunoblot. **e.** Comparison between enrichment values for Effector and Detector components, Lsm proteins and CK subunits in Upf2-TAP purified samples in the presence (grey bars) or absence (blue bars) of *UPF1*. Levels of expression of Upf2 and Upf3 proteins have been verified in this condition in Supplementary Fig. 3.

### Detector forms by binding of a Upf2/Upf3 heterodimer to Upf1 *in vivo*

While the results presented in the previous paragraphs identified a role for Ebs1 and Nmd4 linked with the helicase domain of Upf1, the CH domain of the protein binds the best-known NMD factors Upf2 and Upf3. Among the results that stood out from the purifications of Detector complex components, one of the most striking was the loss of Upf2 and Upf3 in the Upf1 complex after RNase treatment (**Fig. 2a**). This result was unexpected, since a complex can be reconstituted with domains of the three proteins in the absence of RNA (Chamieh *et al*, 2008). The reciprocal experiments using Upf2-TAP and Upf3-TAP as baits showed that Upf1 could be efficiently recovered in the presence or absence of RNase with Upf2-TAP (**Fig. 2b**), but its interaction with Upf3-TAP was sensitive to RNase (**Fig. 2c**). Thus, the role played by RNA in the stability of the purified Upf1-Upf2-Upf3 complex suggested that Detector is assembled on RNA. To investigate the assembly process, we analysed the composition of complexes purified with Upf1-TAP, Upf2-TAP and Upf3-TAP in the absence of *UPF1*, *UPF2* or *UPF3*. Deletion of either *UPF2* or *UPF3* had a major impact on the enrichment of Upf3 and Upf2 with Upf1 (**Fig. 6a, b**). In the absence of *UPF2*, we could no longer detect Upf3 (**Fig. 6a**), which correlates with previously published data that described human Upf2 as bridging Upf1 and Upf3 (Chamieh *et al*, 2008). Unexpectedly, in the absence of Upf3, we could not detect Upf2 in the Upf1-associated complex (**Fig. 6b**). Thus, in the absence of *UPF2*, Upf3 no longer stably associated with Upf1 and in the absence of UPF3, Upf2 was lost from the Upf1-associated factors. We verified that these changes were not due to changes in the total levels of Upf3-TAP and Upf2-TAP in the absence of *UPF2* (**Fig. 6c**) and *UPF3* (**Fig. 6d**). The absence of Upf1 had no impact on the formation of the Upf2-Upf3 complex, independent of which of the two proteins was tagged (**Fig. 6e** and **Supplementary Table 4**). Altogether, these results suggest that Upf2 and Upf3 form a heterodimer independent of Upf1 and that both proteins are required for a stable association with Upf1 in the Detector complex.

### Binding of Effector to NMD substrates depends on Upf2

The formation of Detector on NMD substrates could be followed by a switch to Effector, for RNA decapping and initiation of degradation. Surprisingly, the absence of Upf2 or Upf3 did not alter the protein composition of the purified Effector complex (**Fig. 6a, b**). This puzzling result made us wonder if Effector exists in two forms, one that associates to NMD substrates, and another that would correspond to Upf1 and the decapping machinery in the process of being recycled as RNA-free complexes. To explore this hypothesis we took advantage of the fact that Upf1 binds preferentially to RNAs that are degraded through NMD (Johansson *et al*, 2007). To distinguish between the Effector-bound and Detector-bound Upf1, we purified Effector via Nmd4-TAP (**Fig. 2d**). The 6-fold enrichment of RPL28 pre-mRNA as compared with total RNA in the input fraction (**Fig. 7a**) indicated that a fraction of the Effector complex binds NMD substrates.

**Fig. 7:**
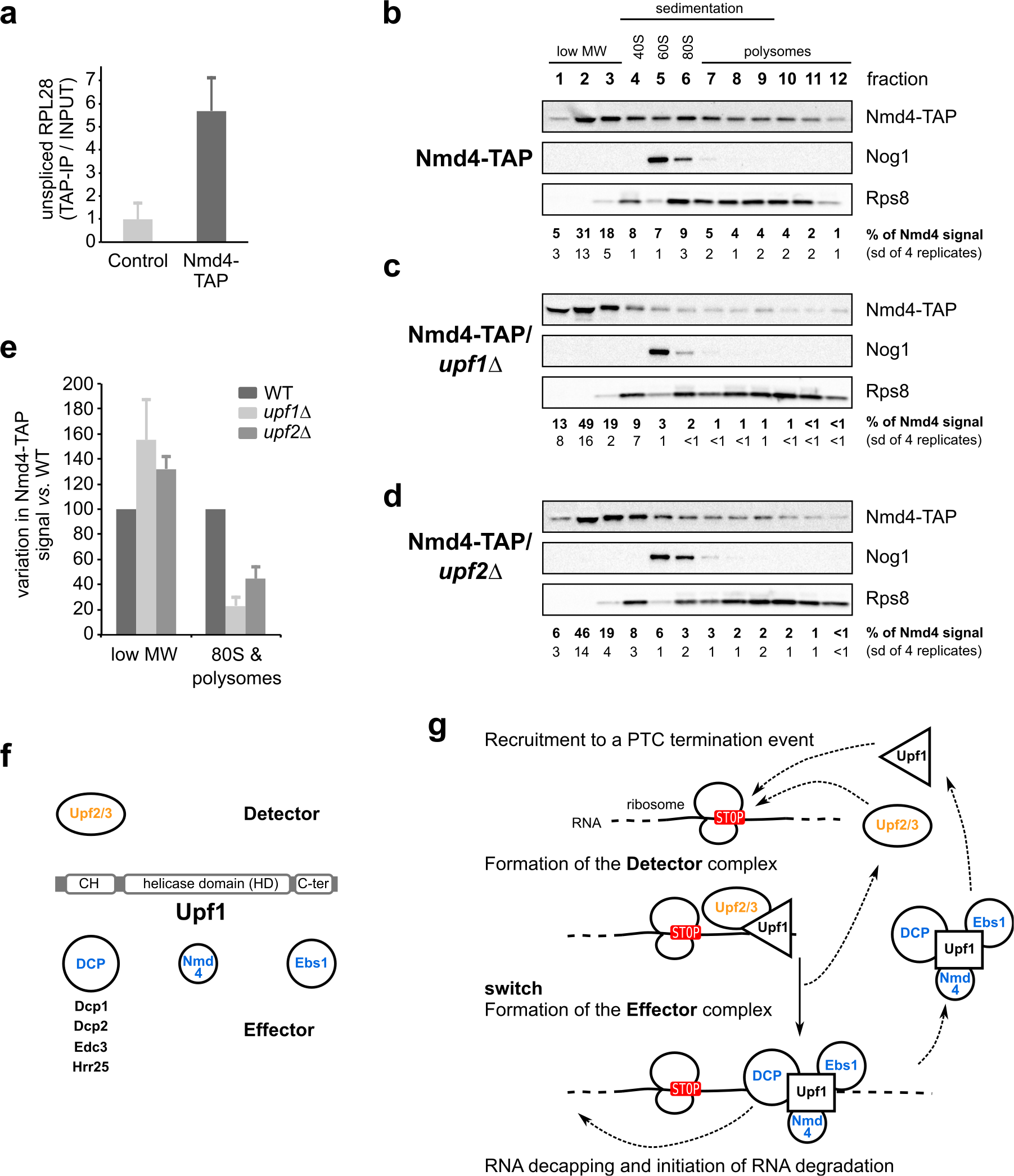
Association of Nmd4 to NMD substrates depends on Detector. **a.** Unspliced RPL28, an NMD substrate, was enriched in Nmd4-TAP purification in comparison with a control untagged strain, as measured by reverse transcription followed by quantitative PCR. Bars correspond to the mean of pre-RPL28 enrichment for 3 independent experiments, as compared with RIM1, a non-NMD mRNA. Error bars correspond to the standard deviation. **b.** to **d.** Distribution of Nmd4-TAP, Nog1 (marker of free 60S subunits) and Rps8 (marker of 40S, 80S and polysome fractions) in wild type ( **b**), *upf1*Δ (**c**) and *upf2*Δ (**d**) strains, tested by immunoblot. Fractions 1, 2, 3 are light fractions, fractions 4, 5 and 6 correspond respectively to ribosomal 40S, 60S and 80S positions, fractions 7 to 12 correspond to polysomes. The percent of total signal for Nmd4-TAP in three independent replicates along with standard deviation of the values are indicated for each fraction. **e.** Quantified relative changes in Nmd4-TAP levels in light and polysomal fractions in *upf1*Δ and *upf2*Δ strains (100% correspond to levels in the wild type strain). **f.** Summary of the Upf1 domains interaction with Detector and Effector components, based on the data presented in this work. **g.** Proposed Detector/Effector sequence of events for NMD in yeast.

We next wanted to know if Effector binding to RNA was dependent on Detector components, Upf2 and Upf3, which would be suggestive of a mandatory sequence of association of the NMD complexes to the RNA substrates. We could not test this hypothesis directly, because deletion of *UPF2* or *UPF3* leads to a massive stabilization of NMD-sensitive RNAs and renders RNA enrichment calculations and comparisons with the wild-type situation unreliable. To overcome this problem, we looked at the association of the Effector complex to RNA by measuring the distribution of Nmd4-TAP in a polysome gradient in the absence or presence of UPF2. While most of Nmd4 sedimented in the upper part of the gradient, a fraction of the protein was found in the polysomes, specifically the monosomal fraction (**Fig. 7b**), which is the fraction that concentrates the majority of NMD substrates (Heyer & Moore, 2016). In the absence of *UPF1*, which mediates all the interactions between Nmd4 and the other components of the effector complex (**Fig. 4b**), Nmd4 was lost from the monosome and polysome fractions (**Fig. 7c**). In the absence of *UPF2*, an essential component of the Detector complex, we also observed a similar change in the distribution of Nmd4 (**Fig. 7d**). Compared with the wild type situation, the absence of Upf1 or Upf2, led to an increase of Nmd4-TAP levels in the lower molecular weight fractions and a decrease in the polysome fractions (**Fig. 7e**). These changes were specific for Nmd4, since the gradient distribution of Rps8, a component of the 40S ribosomal subunits used as a control, was not altered, neither in the *upf1* (**Fig. 7c**) nor in the *upf2*Δ strain (**Fig. 7d**).

The association of Nmd4, a specific component of the Effector complex, to NMD substrates, and its redistribution into lower molecular weight complexes in the absence of the Detector formation, suggest that Detector and Effector are obligatory successive complexes on RNA, and allows us to propose a succession of events compatible with our observations for yeast NMD (**Fig. 7f, g**). The fact that Nmd4 was bound to a region of Upf1 not required for other interactions and still could only be enriched in Effector complexes suggests that Upf1 has a different conformation in Detector and Effector. The switch from a Detector to an Effector conformation described here would permit recruitment of decapping factors, required to degrade NMD substrates (Muhlrad & Parker, 1994), and could be an essential step of the NMD mechanism.

## Discussion

The current study addresses the composition and dynamics of NMD complexes in yeast. We identified two distinct, mutually exclusive (**Fig. 2**) and successive (**Fig. 7**) complexes containing Upf1. We propose that the Detector complex, composed of Upf1, Upf2 and Upf3, is recruited on NMD substrates. A complete re-organization of the initial complex leads to the replacement of the Upf2-3 heterodimer with Nmd4, Ebs1 and the decapping factors Dcp1, Dcp2 and Edc3. Nmd4 and Ebs1 are accessory factors for NMD in yeast and could be functional homologs of human Smg6 and Smg5-7 respectively (**Fig. 4 and 5**). Both the decapping machinery and Upf2-3 interacted with the CH domain of Upf1 (**Fig. 3**), in support of the hypothesis that decapping competes with Upf2-3 to form the Effector. This competition is likely to occur on RNA, since Nmd4 was lost from large RNA-associated complexes in the absence of *UPF2* (**Fig. 7**). Thus, recruitment of decapping on NMD substrates is likely to depend on this switch on RNA, which could be a crucial, albeit not yet explored step in NMD.

How exactly the switch from the “Detector” to an “Effector” complex occurs is unclear, but similar events could be also important for recruiting RNA degradation factors for NMD in other organisms, including humans. Since all the factors that we describe in yeast NMD complexes have mammalian equivalents, and since our results are compatible with a large body of experimental evidence in various other organisms, we propose to use the new global view of NMD to improve and extend the canonical model of NMD (**Supplementary Fig. 7**). We discuss how a revised NMD model, called “Detector/Effector” fits the available experimental data in yeast and other species and to what extent it can serve as a basis to build more complex NMD models that incorporate RNA splicing and Upf1 phosphorylation, two molecular events that do not affect NMD in *S. cerevisiae*.

### Affinity purification strategies for NMD

Before entering into the description of the Detector/Effector model, it is important to see how the results presented here extend and improve those obtained in previous studies. Affinity purification followed by mass-spectrometry is the most effective way to find functional association of proteins and surpasses in precision all the other large-scale interaction methods in yeast (Benschop *et al*, 2010). However, despite the excellent results obtained with this method on hundreds of different complexes, large-scale affinity purification failed to assign specific interactions of Upf1, Upf2 or Upf3 with other yeast proteins (Gavin *et al*, 2006). As a result, most of what was known on the composition of yeast NMD complexes was based on two-hybrid experiments and the use of co-purification and immunoblotting for previously identified factors (e.g. He and Jacobson 1995; Czaplinski et al. 1998; Swisher and Parker 2011). The situation is similar in other species and can be illustrated by the results obtained with the recently published SONAR approach (Brannan *et al*, 2016), which analysed the proteins co-purified with tagged human Upf1 (hUpf1). Human Upf2 was found as the 112th most enriched factor in the list of Upf1-associated proteins and only reached position 28 when the sample was treated with RNase A. Except for Upf3b, ranked second, no other specific NMD factors were identified in the first 100 proteins associated with hUpf1. Thus, without previous knowledge of the identity of NMD factors, the number of false positive results precludes the use of these data to define NMD complexes in human cells.

To understand the molecular mechanisms of NMD, a global and high-resolution view of the composition and dynamics of the involved complexes was missing. This was due to the relative scarcity of some NMD factors (**Fig. 1a**), the dynamic nature of the complexes, their presence in large and heterogeneous RNA-protein assemblies and, potentially, their instability to lengthy incubation periods during purification protocols. To circumvent part of these technical problems, we used a combination of an affinity isolation strategy with an innovative computational workflow. Fast affinity purification preserves transient interactions (Oeffinger *et al*, 2007) and the use of surface-coated magnetic beads gives access to the large RNA-protein complexes in which NMD takes place (Zhang *et al*, 1997) without the bias induced by porous chromatography media (Halbeisen *et al*, 2009). The isolated complexes were analysed by mass-spectrometry in a way that allowed us to extract estimates of the amounts of co-purified proteins. To get rid of abundant contaminants, we used the relative enrichment of proteins (see **Methods**), efficiently highlighting specific interactions.

The top hits obtained in our experiments were validated by immunoblots (**Fig. 2g, 3f**) on co-purified complexes and by second-round purifications by tagging new or known NMD factors and analysing the associated proteins by the same procedure (**Fig. 2**). With these robust quantitative data in hand we discovered novel interactions, including two protein kinases associated with Upf1 (Hrr25; **Fig. 1f, g, 2a** and **Supplementary Table 4**) and Upf2 (Casein Kinase 2 complex; **Fig. 2b** and **Supplementary Table 4**). It also led to the estimation of subtle or drastic changes in the composition of protein complexes that allowed us to test the importance of various NMD factors in the assembly or stability of sub-complexes (**Fig. 4 and 6**). Besides the ability to show the presence of interactions, quantitative mass spectrometry provides a way to define lost or absent proteins in a given complex. This led to the identification of mutually exclusive complexes, very well illustrated by the comparison of Detector components purified via Upf2-TAP with Effector complexes associated with Nmd4-TAP (**Fig. 2b, d**).

The existence of these two distinct complexes in yeast NMD was not known but is compatible with previously published results, like the two-hybrid interactions of Upf1 with Upf2 and Upf3 (He *et al*, 1997), the co-purification of human Dcp1 and Dcp2 with Upf1 (Lykke-Andersen, 2002) or the two-hybrid interactions between yeast Edc3 and Upf1 (Swisher & Parker, 2011). Our results, showing the importance of the CH domain in the interactions between Upf1 and other components of Detector and Effector (**Fig. 3**), are also compatible with previous two-hybrid data showing the importance of this domain in the interactions with Upf2 and Upf3 (He *et al*, 1997) and with Dcp2 (Swisher & Parker, 2011; He & Jacobson, 2015).

Altogether, our enrichment approach and the combination of RNase treated and native complexes analyses drastically reduced the number of NMD-associated candidate proteins to a short list that only contains highly specific top hits and is devoid of factors that are not related to NMD (**Fig. 2h** and **Supplementary Tables 3 and 4**). The extensive measurements of the dynamics of NMD complexes in different mutant conditions bound or not to RNA, support a new hierarchy of the molecular events required for yeast NMD, that we call the Detector/Effector model (**Fig. 7g**).

### A revised universal NMD model

#### Detector formation is both similar and distinct from SURF/DECID

The proposed Detector/Effector yeast mechanism (**Fig. 7g**) can be extended to other species to include elements of the canonical SURF/DECID model that was, until now, restricted to mammalian EJC-enhanced NMD (**Supplementary Fig. 7**). The first step in this revised model involves the recognition of an aberrant translation termination event (Amrani *et al*, 2004; Behm-Ansmant *et al*, 2007) and the formation of a complex containing Upf1, Upf2 and Upf3. Its formation depends on an aberrant translation termination event and could occur through an interaction of Upf2/3 with the translation termination factors eRF1/eRF3 or a terminating ribosome. This type of mechanism has been recently proposed as an alternative to SURF formation in recruiting Upf1 to NMD substrates in human cells (Neu-yilik *et al*, 2017).

Our data support the formation of a Upf1, Upf2, Upf3 complex through the interaction of a Upf2-3 heterodimer (**Fig. 6**) with Upf1 on RNA (**Fig. 2a, c**). The importance of heterodimerisation for the function of Upf2-3 was also observed in *C. elegans,* where co-purification of Upf2/Smg3 with Upf1/Smg2 is strongly reduced in the absence of Upf3/Smg4 (Johns *et al*, 2007). The hypothesis of a recruitment of Upf1 through a Upf2-3 heterodimer is also compatible with the fact that Upf2 and Upf3 co-sediment in yeast polysomal fractions independent on the presence of *UPF1*, while Upf2 looses its association with heavy complexes in the absence of *UPF3* (Atkin *et al*, 1997)

The Detector complex should also contain ribosomal proteins and translation termination factors eRF1 (yeast Sup45) or eRF3 (yeast Sup35). However, since we could not measure an enrichment of these proteins in the complexes purified with Upf1, Upf2 or Upf3 (**Supplementary Table 4**), it is possible that the time of residence of eRF1/eRF3 on stalled ribosomes bound to the Detector complex is short or that the interaction does not resist to the purification conditions. Translation initiation factors, for example, require formaldehyde cross-linking to preserve their association with ribosome initiation complexes (Valásek *et al*, 2007). The instability of the interaction between termination factors and NMD complexes is also suggested by the fact that the association between human Upf1 and eRF3 could not be detected in complexes purified with chromosomal tagged Upf1-TAP (Schell *et al*, 2003). The presence of eRF3 in NMD complexes was only observed with a C126S variant of human Upf1, which is no longer competent for NMD (Kashima *et al*, 2006), and when FLAG-Upf1 was overexpressed (Hug & Cáceres, 2014).

#### Smg1 phosphorylation step and links with other Smg factors

While the formation of the Detector complex is the first step in the revised NMD model, the phosphorylation of the SQ-rich C-terminal region of Upf1 by the protein kinase Smg1 can be incorporated as an optional step. Depending on the species, phosphorylation could be essential, facultative or completely absent. However, we show here that the presence of the C-terminal region of Upf1 remains important for NMD in yeast, as demonstrated by its effect on the ability of Upf1 fragments to complement the deletion of the gene (**Fig. 5f**) and by its role in binding Ebs1 (**Fig. 3e**). Thus, the C-terminal extension of yeast Upf1 seem function like the SQ C-terminal extension in other species, even in the absence of the characteristic SQ/TQ motifs and phosphorylation. The C-terminal extension of yeast Upf1 can thus be the functional equivalent of the C-terminal Upf1 extension in other species.

Smg1 protein equivalents are associated with the phosphorylation of the Upf1 C-terminal SQ-rich extension (Page *et al*, 1999; Yamashita *et al*, 2001; Okada-Katsuhata *et al*, 2012). This phosphorylation is crucial for NMD efficiency in several species and phosphorylated residues are important to recruit Smg5-7, to which Ebs1 is similar (**Fig. 5a**) (Luke *et al*, 2007). Smg1 phylogeny indicates that the kinase was present in the last eukaryotic common ancestor (LECA) but has been independently lost in fungi, *A. thaliana* but not other green plants, red and brown algae, excavates such as *Trypanosoma brucei* and *Giardia lamblia* (Lloyd & Davies, 2013) and ciliates, such as *Tetrahymena thermophila* (Tian *et al*, 2017). However, these species present active NMD mechanisms that depend on equivalents of Upf1, Upf2 and Upf3, suggesting that phosphorylation of the C-terminal region is not an absolute requirement for NMD in these species. In line with this observation, no phosphorylated residues have been reported in yeast Upf1 C-terminal region, even if phosphorylated residues have been detected in Upf1 (Lasalde *et al*, 2013). These observations indicate that the C-terminal domain *per se* is important for NMD and that the main role of the C-terminal extension, with or without phosphorylation, is the recruitment of later factors and the formation of Effector complexes. In support to this idea, a chimeric Upf1 where the helicase domain of the yeast protein was replaced with the similar region of human Upf1 partially complements the deletion of *UPF1* in yeast as long as it preserves the fungal-specific N and C-terminal extensions (Perlick *et al*, 1996).

The similarity between the binding of Ebs1 to the C-terminal domain of yeast Upf1 and the recruitment of Smg5/7 to phosphorylated SQ residues in metazoan NMD also extends to Nmd4. Nmd4 is a specific marker of the Effector complex that directly binds the helicase domain of Upf1 (**Fig. 4**) and consists of a PIN domain similar with the PIN domain of mammalian Smg6 (**Fig. 5a**). Nmd4 binding to the helicase domain of yeast Upf1 correlates with the known phosphorylation-independent binding of Smg6 to a region of Upf1 helicase domain (Chakrabarti *et al*, 2014; Nicholson *et al*, 2014). Thus, the architecture and function of Upf1 and the associated factors seem to be more conserved than previously thought.

#### EJC-independent and EJC-enhanced NMD are two versions of the same mechanism

One of the major benefits of the revised NMD model is to provide a single mechanism for EJC-independent and EJC-enhanced RNA degradation. For EJC-enhanced NMD, the interaction of Upf2-Upf3 with EJCs (Lykke-Andersen *et al*, 2001; Le Hir *et al*, 2001) deposited downstream aberrant termination codons could ensure, for example higher local concentrations of the factors and stimulate the initial steps in the formation of a Detector complex. The variations in the importance of EJCs for NMD in various organisms and the fact that physiologic substrates of NMD, such as GADD45A, are degraded through an EJC-independent NMD mechanism (Chapin *et al*, 2014), further support the idea that EJC-enhanced and EJC-independent RNA degradation are alternatives of the same fundamental molecular mechanism.

#### A molecular switch around Upf1 for the Detector to Effector transition

The defining feature of the revised model is the complete change in composition of Upf1-bound complexes from the Detector to the Effector. Upf1 conformation is likely to be dramatically different between the two complexes since: Upf2 and Upf3 were completely absent from complexes purified using Nmd4-TAP (**Fig. 2d**); Nmd4 was bound to a region of Upf1 that is different from the N-terminal CH domain, to which decapping factors and Upf2-3 bind (**Fig. 3c, d**); structures of Upf1 with and without a Upf2 C-terminal fragment show a different organisation of the Upf1 helicase sub-domains 1B and 1C (Clerici *et al*, 2009); all the interactions of Nmd4 with Effector components depend on Upf1 (**Fig. 4b**). This change was not apparent in previous studies that lacked global quantitative estimations of all the components of purified complexes at the same time.

The main result of the switch is the recruitment of the decapping enzyme, specifically the core decapping proteins Dcp1, Dcp2 and Edc3 (**Fig. 2a, d, g**). Decapping is the first and major step in the degradation of NMD substrates in yeast (Muhlrad & Parker, 1994) and is responsible for the degradation of about a third of NMD substrates in human cells (Lykke-Andersen *et al*, 2014). The other two-thirds of NMD substrates in human are likely to be degraded through a pathway that depends on the presence of catalytically active Smg6 (Eberle *et al*, 2009; Huntzinger *et al*, 2008). In view of the modest increase in NMD substrates levels in an *nmd4Δ* mutant (**Fig. 5** and **Supplementary Fig. 5**), it is unlikely that Nmd4 PIN domain has an endonucleolytic activity in yeast NMD. However, its tight association with Upf1 **(Fig. 2a, d** and **4d**), and its major effect on NMD triggered by overexpression of a truncated Upf1 (**Fig. 5f**) suggest that Nmd4 is a Upf1 co-factor that assists the helicase in yeast NMD. The importance of Nmd4 in fungal NMD is supported by the presence of orthologs in many species, including *Yarrowia lipolytica* (Pryszcz *et al*, 2011), yeast that is as distant from *S. cerevisiae* as the urochordate *Ciona intestinalis* is from humans (Dujon *et al*, 2004). Thus, it is likely that different organisms use the same basic molecular machineries to degrade NMD substrates, but the balance between decapping, endonucleolysis or other alternative degradation processes is variable depending on each species.

#### Conclusion

Our data and the extended Detector/Effector model provides a solution to the long-term controversy about the conservation of NMD molecular mechanisms among eukaryotes and will allow future work on yet unsolved issues: how aberrant termination is recognized and leads to Detector formation, where on RNA are positioned Detector and Effector complexes, how Effector and Detector components are recycled and at what steps ATP binding and hydrolysis by Upf1 play crucial roles.

## Author Contributions

M.D., A.N. and C.S. conceived the project. M.D., L.D., A.N., C.P., J.K., H.L-H., A.J., and C.S. designed experiments. M.D., L.D., A.N., J.K, and C.S. performed experiments. A.N., L.D, M.D., and C.S. analyzed the data. M.D. and C.S. wrote the manuscript. All authors discussed results and participated to the corrections of the manuscript.

## Acknowledgments

We thank Magalie Duchateau, Julia Chamot-Rooke and Mariette Matondo of the Mass Spectrometry for Biology UtechS at the Pasteur Institute for access to the mass-spectrometry facility, Jean-Yves Coppée, from the Transcriptome and Epigenome, Biomics platform for access to the sequencing facility. We thank Giorgio Dieci, University of Parma, Italy, for the Rps8 antibodies. We thank members of the Genetics of Macromolecular Interactions Unit: Gwenaël Badis for providing RNA sequencing results for the *upf1*Δ strain, helpful discussions and critical reading of the manuscript, Antonia Doyen for performing initial affinity purification experiments and for building strains, Frank Feuerbach for providing strains, reagents, critical suggestions, advice and support in writing the manuscript, and Micheline Fromont-Racine for discussions and critical reading of the manuscript. We thank Christophe Malabat from the Bioinformatics and Biostatistics hub, Pasteur Institute, for help with data analysis and support in making the results available through a web interface. The study was supported by the ANR CLEANMD grant (ANR-14-CE10-0014) from the French Agence Nationale de la Recherche to C.S., A.J. and H.L-H. and by the Ministère de l’Enseignement Supérieur et de la Recherche and the “Fondation ARC pour la recherche sur le cancer” PhD fellowships to M.D., and by continuous financial support from the Institut Pasteur and Centre National de Recherche Scientifique, France.

## Supplementary Information

### Contents

#### Supplementary Figures

Fig. S1 – Workflow for quantitative analysis of MS/MS results from affinity purified complexes.

Fig. S2 – N-terminal and C-terminal tagged Upf1 enrich similar sets of specific proteins.

Fig. S3 – Controls of total tagged protein levels in the presence or absence of other NMD components.

Fig. S4 – Alignment of hSmg6, hSmg5, hSmg7, Ebs1 and Nmd4 domains sequences.

Fig. S5 – Deletion of NMD4 and EBS1 stabilize a set of transcripts that is also stabilized in the absence of UPF1.

Fig. S6 – The helicase domain of Upf1 alone can destabilize RPL28 pre-mRNA, an NMD reporter.

Fig. S7 – Comparison between the canonical SURF/DECID model and our extended Detector/Effector model.

#### Methods

Contact for reagent and resource sharing.

Experimental model and subject details.

Method details.

Quantification and statistical analysis.

Data and software availability.

References.

#### Strains, oligonucleotides and plasmid tables

##### Supplementary Tables

Table **S1** – LTOP2 values for the proteins identified in affinity purified samples

Table **S2** – Average LTOP2 values and standard deviation of the results from independent replicate experiments.

Table **S3** – Enrichment values in purification over total protein content (log2 transformed values).

Table **S4** – Average enrichment values and standard deviation of the results from independent replicate experiments.

Table **S5** – Summary of the number of replicates, the number of proteins robustly quantified and the efficiency of RNase treatment for each purification type.

Table **S6** – RNASeq raw data analysis results summary.

Note: tables **S1** to **S4** are provided as a single separate Excel Office Open XML file, and **S5** and **S6** as individual spreadsheet files.

**Fig. S1.**
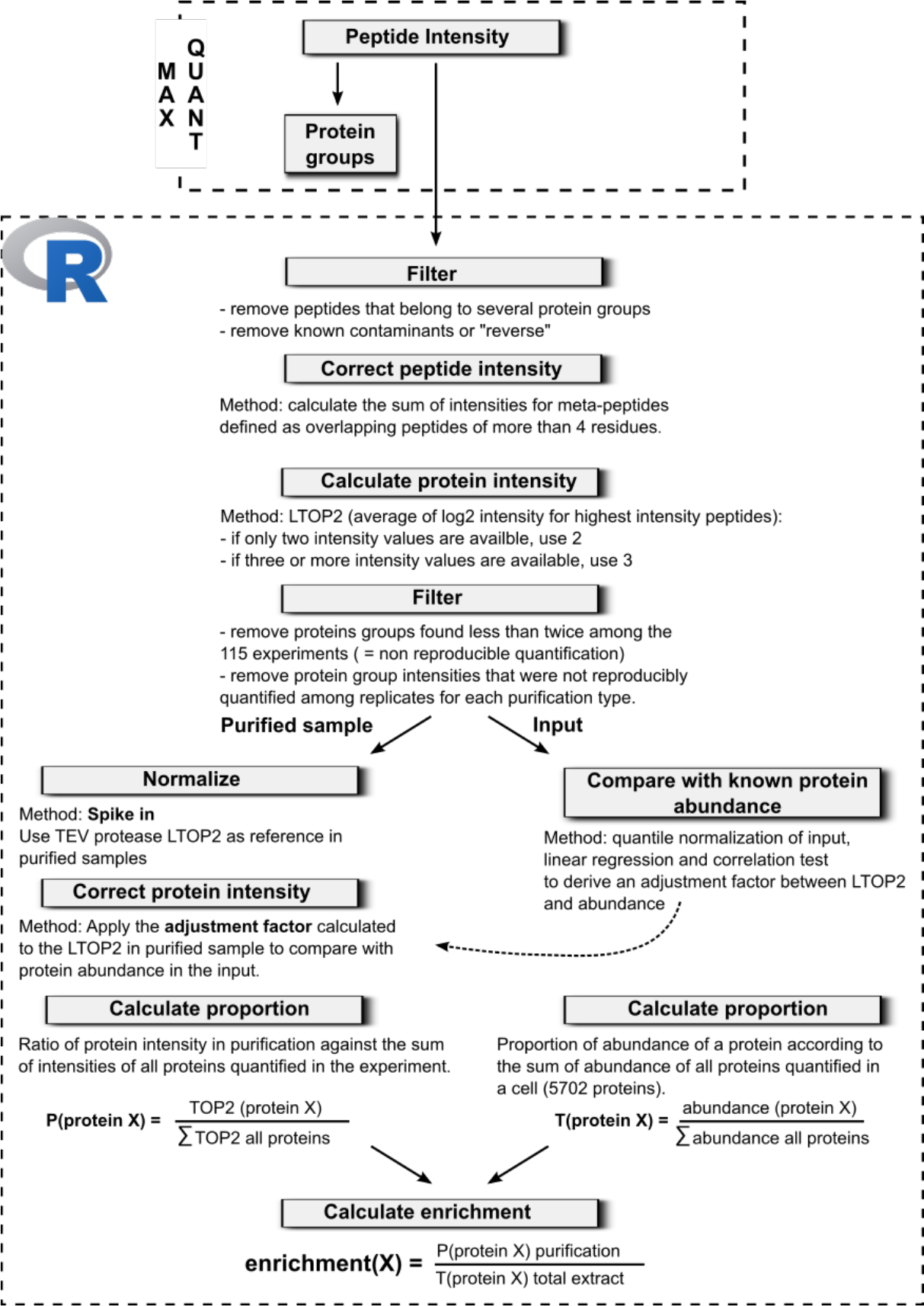
Workflow for quantitative analysis of MS/MS results from affinity purified complexes. Notes about the depicted procedure:

1. MaxQuant output for peptide intensities and their association with a given protein was used as the main input for computing LTOP2 scores and enrichment. To calculate the false discovery rate (FDR) of the MS/MS analysis, MaxQuant builds reverse sequence « artificial » proteins that serve as negative controls for the identification procedure. Reversed sequences and common contaminants (trypsin, keratins) were removed from further analyses in the early steps of the analysis.
2. A protein group corresponds to a single protein or several proteins with very high sequence similarity that cannot be discriminated by peptide analysis. For further analyses, we used the identity of the protein of the group with most coverage.
3. The TEV protease was added in each purification experiments with the same relative amount to elute complexes from beads. This step is important to be able to compare replicates between them and the different purification types.
4. The comparison of our input LTOP2 with the abundance data from Ho et al. 2017, allowed to calculate a factor that served to adjust LTOP2 values and make them compatible with the dynamic range and scale of published abundance values.

**Fig. S2.**
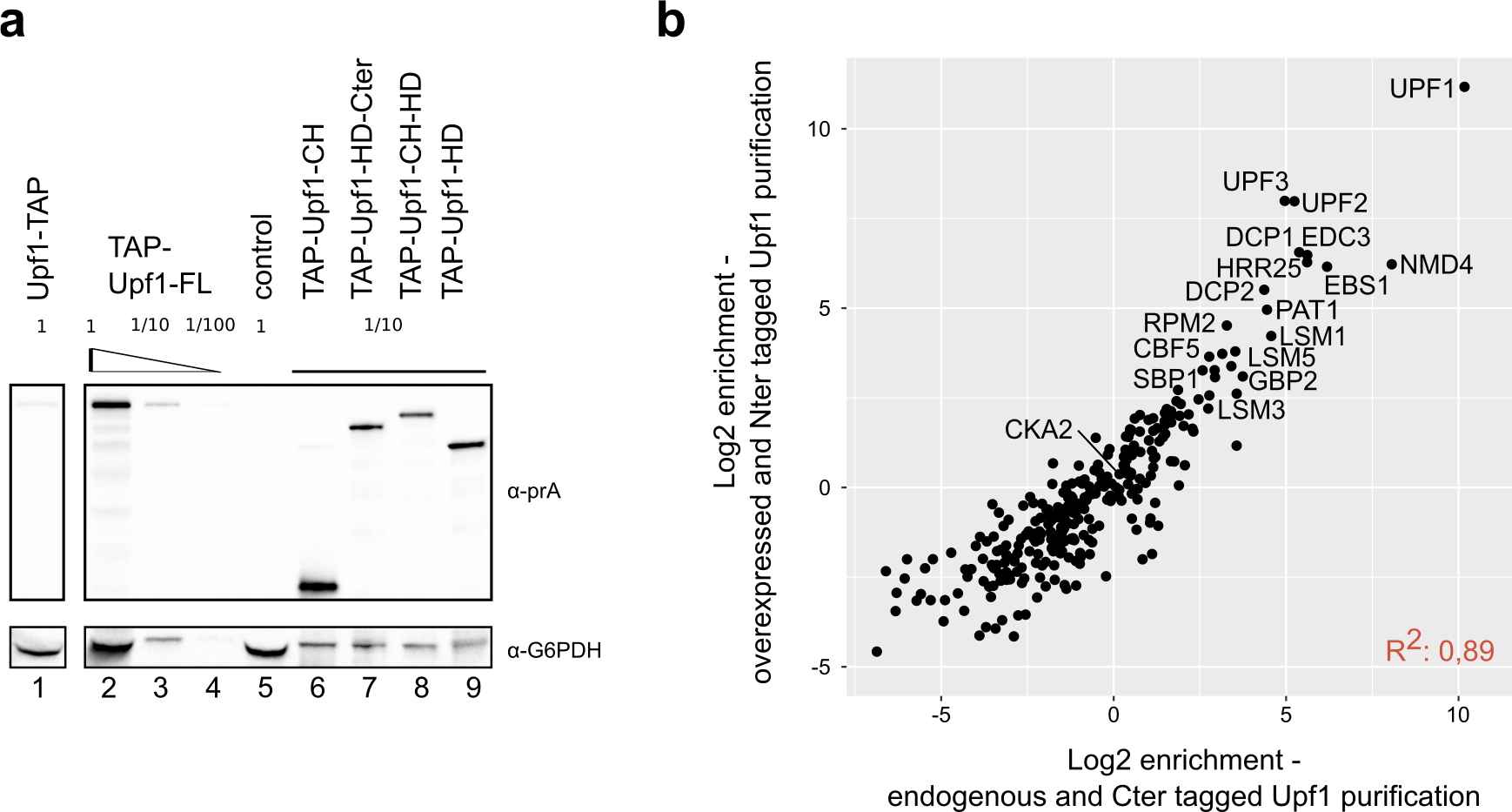
N-terminal and C-terminal tagged Upf1 enrich similar sets of specific proteins. **a)** Estimation of the levels of overexpression for N-terminal tagged Upf1 fragments, in comparison with chromosomally C-terminal tagged protein. G6PDH was used as a loading control. Serial dilutions were used to test the ability of the immunoblot signal to estimate protein levels. **b)** Enrichment values for purifications done with chromosomally C-terminal tagged Upf1 (x axis) and N-terminally TAP tagged Upf1.

**Fig. S3.**
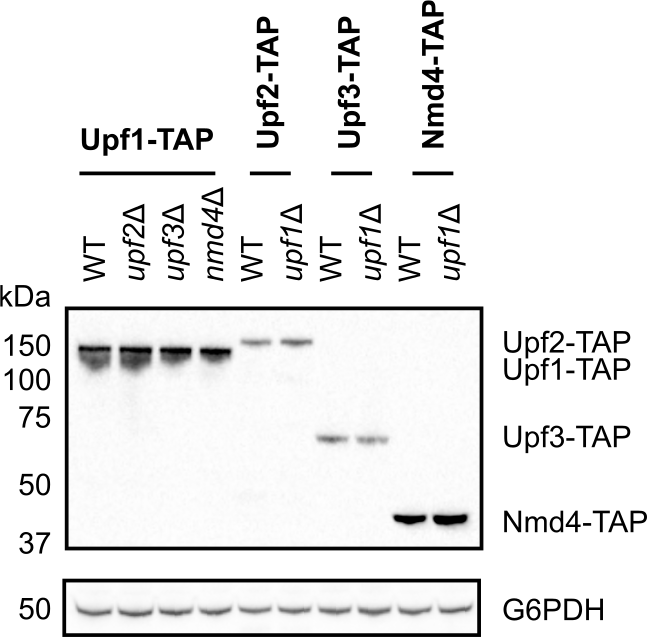
Controls of total tagged protein levels in the presence or absence of other NMD components. Total protein extracts from the described strains were tested by immunoblot to detect the protein A part of the TAP tag. G6PDH was used as a loading control.

**Fig. S4.**
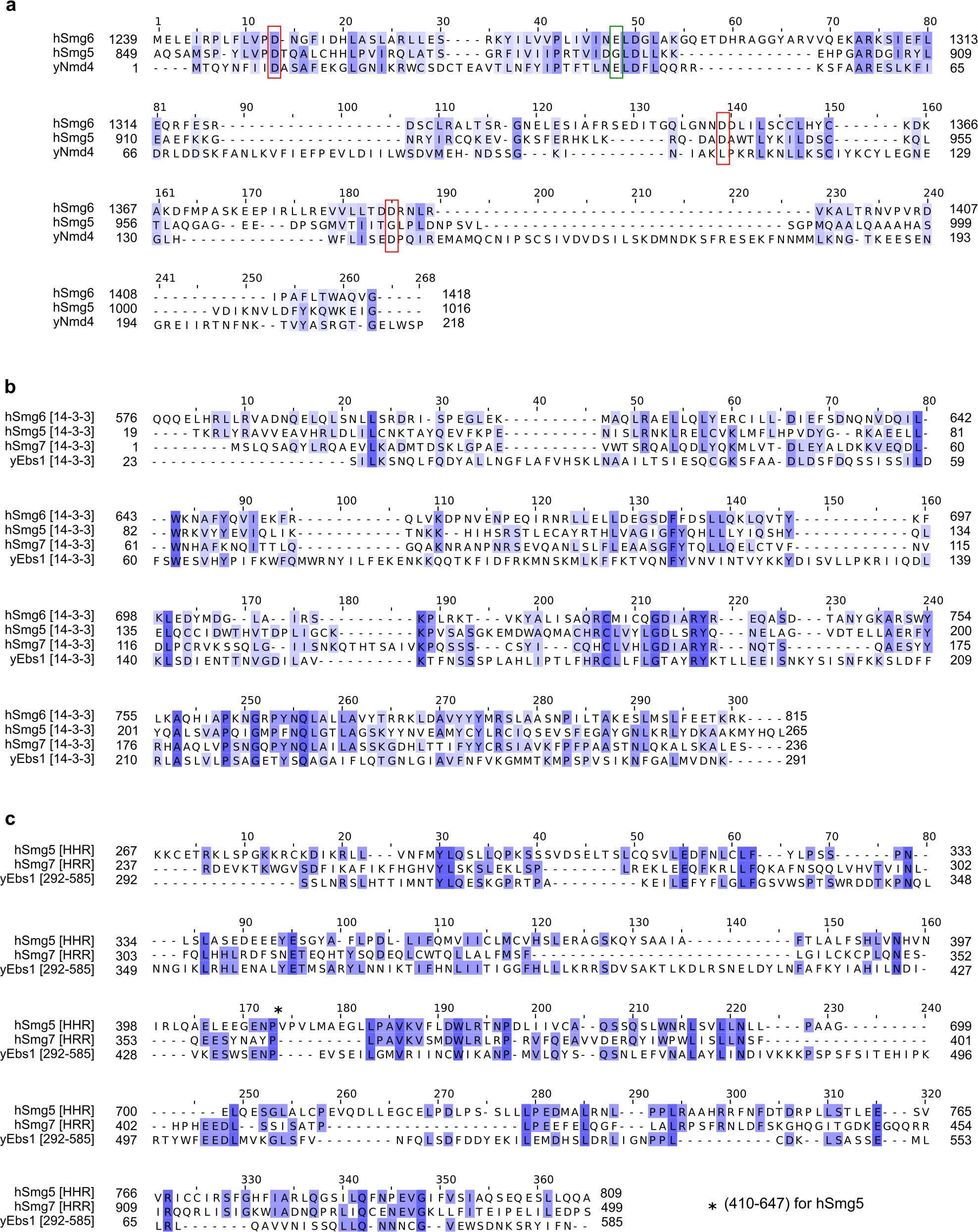
Alignment of hSmg6, hSmg5, hSmg7, Ebs1 and Nmd4 domains sequences. Alignment of the PIN domains of hSmg6, hSmg5 and ScNmd4 **(a)**, the 14-3-3 domains of hSmg6, hSmg5, hSmg7 and Ebs1 **(b)** and of the helical hairpin region (HHR) of hSmg5, hSmg7 and Ebs1 (292-585) **(c)**. These alignments were obtained with the algorithm Mafft with default parameters; colour represents percentage of identity or similarity (BLOSUM62).

**Fig. S5.**
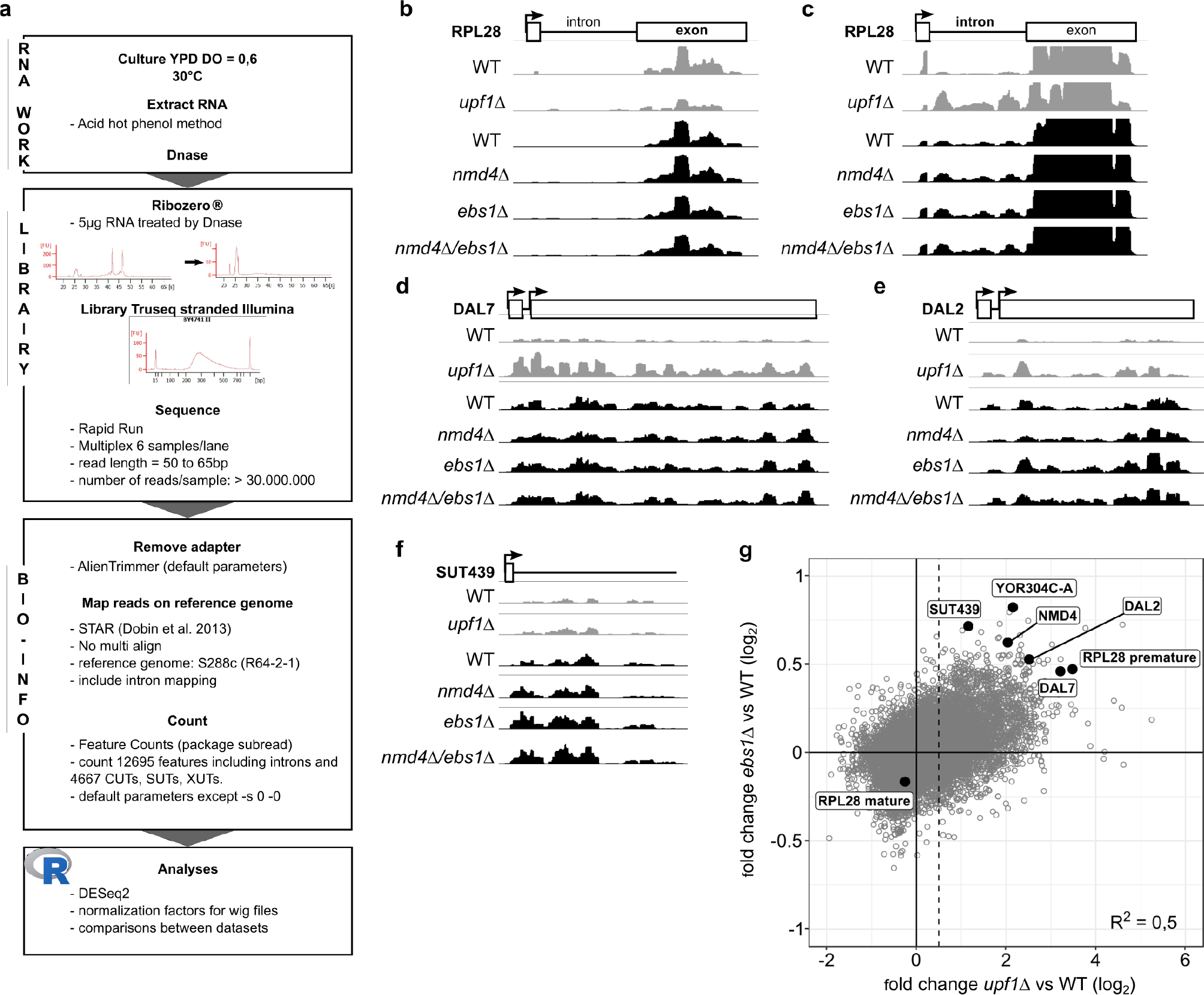
Deletion of *NMD4* and *EBS1* stabilize a set of transcripts that is also stabilized in the absence of *UPF1*. **a)** Workflow used for RNAseq experiments and analyses. (B) to (G) Examples of NMD substrates sequencing profiles in WT, *upf1*Δ, *nmd4*Δ and *ebs1*Δ experiments. NMD substrates belong to different classes, intron containing (RPL28; **b** and **c**), uORF (DAL7, DAL2; **d** and **e**), non-coding RNA (SUT439; **f**) and long 3’UTR (YOR304C-A; **g**). Profiles were normalized using the samples median counts. For RPL28 (**b** and **c**), we represented the profile of the entire transcript showing that the signal in the exon is similar in each strain (**b**). A zoom of the intron region (**c**) shows a higher signal in this specific region of unspliced RPL28 in mutants by comparison of WT. H. Scatter plot of transcript log2 fold change in *upf1*Δ against *ebs1*Δ. The dashed line represents the limit over which RNA are considered as stabilized, a value of 0.5 in log2 (1.4 fold change).

**Fig. S6:**
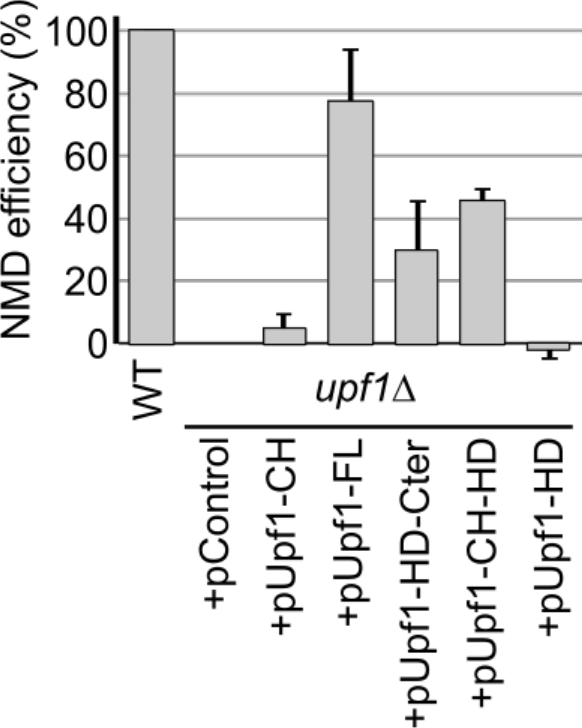
The helicase domain of Upf1 alone can destabilize RPL28 pre-mRNA, an NMD reporter. Total RNA from wild-type or *upf1*Δ strain transformed with an empty plasmid (pControl) or plasmids expressing various Upf1 fragments (see Figure 3A) was tested by reverse transcription and quantitative PCR. The levels of RPL28 pre-mRNA were normalized using an NMD-insensitive transcript (RIM1) and an NMD efficiency score was calculated based on the difference between a wild type and a *upf1*Δ strain.

**Fig. S7:**
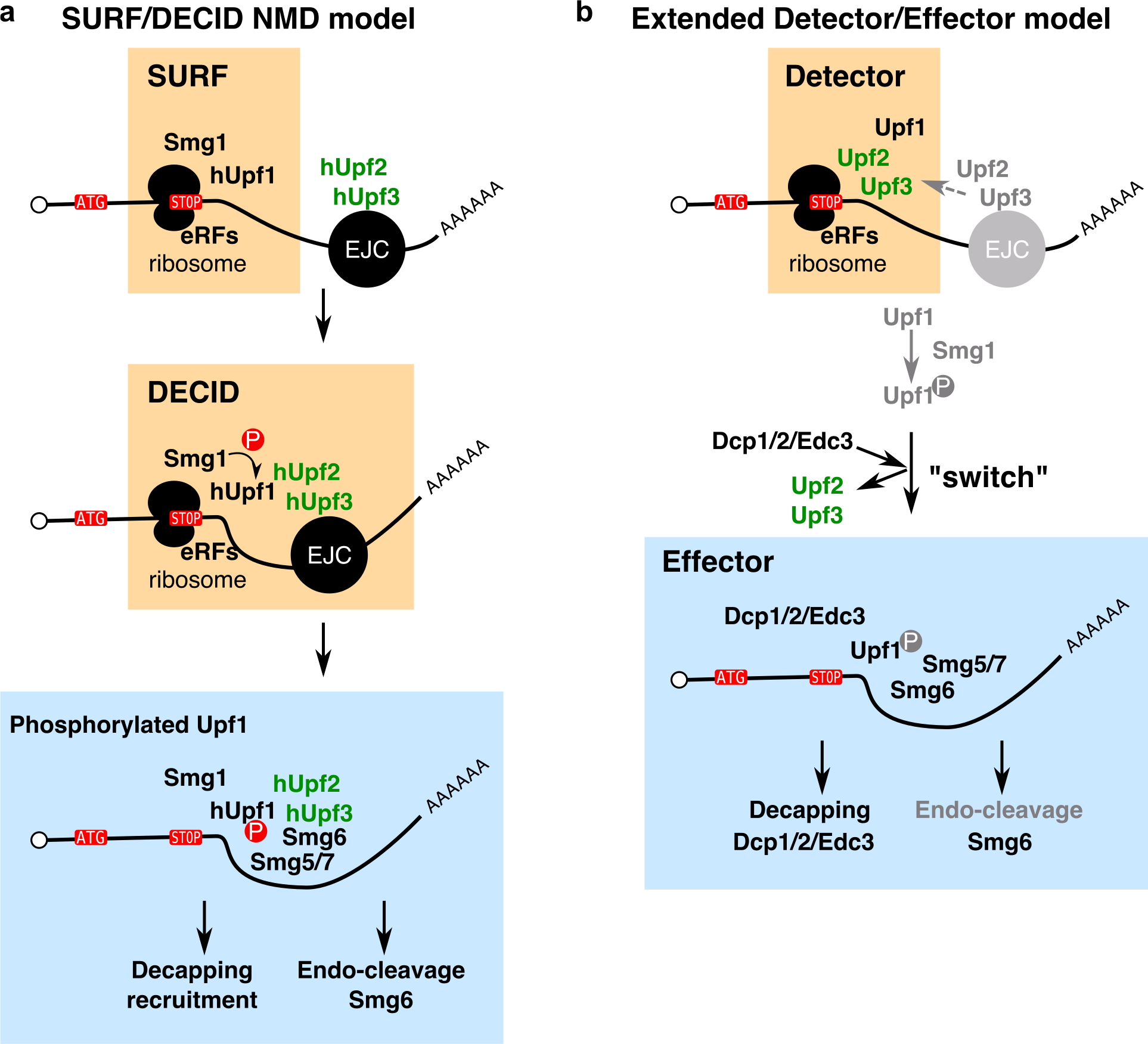
Comparison between the canonical SURF/DECID model **(a)** and our extended Detector/Effector model **(b)** for NMD. Orange and blue squares mark equivalent steps in both models. Light grey elements in the revised model represent optional steps that can further enhance the NMD process under certain conditions and in specific organisms.

## EXPERIMENTAL MODEL AND SUBJECT DETAILS

### Yeast strains

*Saccharomyces cerevisiae* strains were derived from BY4741 (mat a) and BY4742 (mat alpha) strains. The full list of strains is provided as a table in the Supplementary material.

### Bacterial strains

*E. coli* strain NEB 10-beta (NEB Cat# C3019) was used for construction of plasmids by Gibson assembly and their multiplication. *E. coli* BL21(DE3) cells were used for expression of recombinant Nmd4 and Upf1 proteins.

## METHOD DETAILS

### Media and growth conditions

Yeast cells were grown in YPD (20g·L^−1^ glucose, 10g·L^−1^ yeast extract, 20g·L^−1^ bacto-peptone, 20g·L^−1^ bacto-agar for plates only) and in synthetic media without uracil to select transformants and maintain URA3 plasmids. All yeast strains were freshly thawed from frozen stocks and grown at 30°C.

Bacterial strains were grown in LB media, supplemented with antibiotics when necessary, at 37°C.

### Yeast strains construction

Yeast strains used in this paper are listed in a specific table. C-terminal TAP-tagged strains originated from the collection of systematically built strain (Ghaemmaghami *et al*, 2003). For some of the strains, we have modified the classical TAP tag (CBP-TEV cleavage site-protein A) and added an additional 6-His tag between CBP and the TEV cleavage site. To do that we have used the pCRBlunt-CRAP(6-HisTAP) plasmid that can replace the original tag by homologous recombination. TAP or CRAP (CBP-6His-TEV-proteinA) tagged versions of proteins were used for our experiments and gave similar results; both versions bear the “TAP” name in the text, as only the protein A part of the tag and the TEV cleavage site were used in our experiments.

Deletion strains were part of the systematic yeast gene deletion collection (Giaever *et al*, 2002) distributed by EuroScarf (http://www.euroscarf.de/) or were built by transformation of BY4741 strain with a cassette containing a selection marker cassette flanked by long recombination arms located upstream and downstream the open reading frame of the gene. Deletions were tested by PCR amplification of the modified locus.

### Plasmids construction

N-terminal TAP-tagged Upf1 fragments were expressed from single copy plasmids derived from pCM189 (Garí *et al*, 1997) under the control of a Tet-OFF promoter. Cloning of Upf1 fragments into plasmid was done by a modified version of Gibson assembly (Fu *et al*, 2014) using NotI digested pCM189-NTAP (plasmid 1233, Saveanu *et al*, 2007) and the PCR product obtained with oligonucleotides CS1359 and CS1364 (UPF1-FL), CS1359 and CS1361 (UPF1-CH), CS1362 and CS1364 (UPF1-Cter), CS1359 and CS1393 (UPF1-NoCex) or CS1362 and CS1393 (UPF1-Cter-NoCex) on template pDEST14-NAM7 (pl. 1350) or pDONR201-NAM7 (pl. 1330). Briefly, we mixed 30 to 50ng of linearized vector with 60 to 300ng of amplified PCR insert(s) in 200μL microtubes containing 10μL of 2x Hot Fusion Buffer: 2x pre-assembly buffer 5x (0.5M Tris pH 7.5, 50mM MgCl2, 1mM each dNTP, 50mM DTT, 25% PEG-8000) with 0.0075u·μL-1 of T5 exonuclease and 0.05u·μL-1 of Phusion Hot Start DNA polymerase. Distilled water was added to the tube for a final volume of 20μL. Tubes were incubated in a thermocycler for 1 hour at 50°C, then slowly ramped down to 20°C in 5 minutes (0.1°C per second), and held at 10°C. The Hot Fusion reaction was used for transformation or stored at −20°C if not used immediately. Otherwise, 1μL of the reaction was transformed in *E. coli* NEB 10-beta competent cells. Insertion of *UPF1* fragments into vector was verified by restriction enzyme digestion and sequencing.

Coding sequences of yeast full length *NMD4*, yeast *UPF1* helicase domain, and human *UPF1* helicase domains (Uniprot accession codes Q12129, P30771, and Q92900-2 respectively) were cloned between NheI/NotI, NheI/XhoI, and NdeI/XhoI in variants of pET28a (Novagen). In these vectors the NcoI– NdeI cassette was either deleted by mutating the NcoI site to an NdeI site or replaced by the coding sequence for a His-tag or a CBP tag followed by a TEV protease cleavage site. In the vector without the NcoI-NdeI cassette, a TEV protease cleavage site was engineered in front of the C-terminal His-tag. Fragments of human *UPF1* (helicase domain, 195-914), yeast *UPF1* (helicase domain, residues 220–851), and *NMD4* (Full length, residues 1–883) were amplified by PCR with the appropriate restriction sites, using oligo pairs HLH725/HLH726, HLH2705/HLH2706 and MD99/MD100 respectively. PCR products were purified using PCR cleanup Qiaquick (*NMD4*) or Promega (human and yeast *UPF1*) kits, digested for 1 hour at 37°C using NEB Cutsmart buffer and the corresponding enzyme couples, then further gel purified using Quiaquick kit (*NMD4*) or Promega gel purification kit (human and yeast *UPF1*). Plasmids pHL5 and pHL4 were digested in parallel using NheI/NotI (pHL5), NheI/XhoI (pHL4) or NdeI/XhoI (pHL4) and purified on 0.8% agarose gels. Digested PCR products were mixed with corresponding plasmids in a 1:3 molar ratio and ligated overnight at 16°C in a 15μl reaction using T4 DNA ligase. Ligase was heat-inactivated 10 minutes at 65°C, then ligation products were used to transform *E. coli* NEB 5-alpha competent cells. Insertion of *NMD4* and *UPF1* fragments into vectors was verified by restriction enzyme digestion and sequencing.

### Total protein extracts and western blots

Total protein extracts were prepared from 5 *A*_600_ of exponential culture with a fast method using alkaline treatment (Kushnirov, 2000). Cells were incubated in 200μL of 0.1M NaOH for 5 min at room temperature, collected by 3 min centrifugation and resuspended in 50μL of sample buffer containing DTT (0.1M). Proteins were denatured for 3 min at 95°C, and cellular debris were pelleted by centrifugation. 10μL of supernatant or diluted supernatant (for quantification scale) were loaded on acrylamide NuPAGE Novex 4-12% Bis-Tris gels (Life technologies). After transfer to nitrocellulose membrane with a semi-dry fast system (Biorad trans-blot) with discontinuous buffer (BioRad technote 2134), proteins were detected by hybridization with appropriate antibodies.

### TAP affinity purifications

TAP-tagged proteins were purified using a one step purification method. Briefly, frozen cell pellets of 2 to 8L culture were resuspended in lysis buffer (20mM Hepes K pH7.4, 145mM KCl, Protease inhibitor, 1X Vanadyl-Ribonucleoside Complex, 40u·mL^−1^RNasin) and lysed using a French press (2 passages at 1200 PSI). The lysate was cleared at 4°C for 20min in a JA-25.50 rotor (Beckman) at 15 000rpm. Magnetic beads (Dynabeads M-270 epoxy) coupled to IgG were added to the protein extract and incubated for 1 hour at 4° (Oeffinger *et al*, 2007). Beads were collected with a magnet and extensively washed three times with a first buffer (HKI: 20mM Hepes K pH7.4, 145mM KCl, 40u·mL^−1^ RNasin, 0.1% Igepal) and twice with a second buffer (HK: 20mM Hepes K pH7.4, 145mM KCl, 40u·mL^−1^RNasin, 1mM DTT). The lack of detergent in the second buffer ensured compatibility with mass spectrometry. After washing, complexes were eluted by incubation in elution buffer (HK + recombinant purified AcTEV, Thermo Fisher Scientific) for 1 hour at 16°C on a rotating wheel. Supernatant that contained purified complexes was collected and precipitated using the methanol/chloroform method (Wessel & Flügge, 1984) before further analysis by silver staining, western blot or mass spectrometry.

In case of RNase treatment, lysed cells were divided in two equal parts, one part was treated with RNase A and RNase T1 and the other part was not. Purifications were done in parallel. For treated samples, clarified lysate were treated a first time with 300u·mL^−1^ of RNase T1 (Thermo scientific, Cat#ENO542) and 167.10^5^ u·mL^−1^ of RNase A (Thermo scientific, Cat#ENO531), 20 minutes on ice. The sample was also treated a second time, after adsorption of proteins on beads and washing, using a first wash with HK buffer with RNase T1 at 83u·mL^−1^ of the clarified lysate previously used and RNase A at 17u·mL^−1^ of the clarified lysate previously used for 20 minutes at 25°C.

### Affinity purification for RNA analysis

We used 4L yeast culture at exponential phase, *A*_*600*_ 0.6 to 0.8 and processed with the same method as TAP immuno-purification but with more precaution to work fast, on ice and in an RNase free environment. In addition, buffers were freshly prepared, lysis buffer contained 5mM MgCl2 to maintain mRNP integrity, washing steps were reduced to three washes with HKI + DTT buffer and elution was done in HKI + DTT + AcTEV. After elution, RNA extraction was done by 3 steps of acid phenol/chloroform followed by overnight ammonium acetate precipitation at-20°C. RNA pellet were resuspended in water and processed for DNase treatment (Ambion TURBO DNA-free kit) and RT-qPCR analysis.

At the first RNA extraction step, we also collected proteins that were at the interface between aqueous phase (RNA) and organic phase. Proteins were precipitated by adding 100% ethanol, incubating at-20°C for 1h and centrifugation. Washed and dried protein pellets were used for western blotting to verify purification efficiency.

### Mass spectrometry acquisition and data analysis

Protein samples were treated with Endoprotease Lys-C (Wako) and Trypsin (Trypsin Gold Mass Spec Grade; Promega). Peptide samples were desalted using OMIX C18 pipette tips (Agilent Technologies). The peptides mixtures were analyzed by nano-LC-MS/MS using an Ultimate 3000 system (Thermo Fisher Scientific) coupled to an LTQ-Orbitrap Velos mass spectrometer or LTQ-Orbitrap XL (Thermo Fisher Scientific). Peptides were desalted on-line using a trap column (C18 Pepmap100, 5μm, 300μm×5mm (Thermo Scientific)) and then separated using 120min RP gradient (5–45% acetonitrile/0.1% formic acid) on an Acclaim PepMap100 analytical column (C18, 3μm, 100Å, 75μm id x 150mm, Thermo Scientific) with a flow rate of 0.340μL·min^−1^. The mass spectrometer was operated in standard data dependent acquisition mode controlled by Xcalibur 2.2. The instrument was operated with a cycle of one MS (in the Orbitrap) acquired at a resolution of 60,000 at m/z 400, with the top 20 most abundant multiply-charged (2+ and higher) ions subjected to CID fragmentation in the linear ion trap. An FTMS target value of 1e6 and an ion trap MSn target value of 10000 were used. Dynamic exclusion was enabled with a repeat duration of 30s with an exclusion list of 500 and exclusion duration of 60s. Lock mass of 445.12002 was enabled for all experiments.

### Protein expression and purification from *E. coli*

Plasmids pHL1484 (CBP-*NMD4* full length-6His), pHL1301 (yeast *UPF1* helicase domain-6His) and pHL201 (human *UPF1* helicase domain-6His) were used to transform BL21 (DE3) Codon Plus competent cells. After plating and overnight cell growth on Kanamycin LB plates, 3-4 colonies were inoculated in 25 ml LB media supplemented with Kanamycine (50mg.L^−1^). After 6 hours of incubation at 37°C, each starter culture was used to inoculate a 1L LB culture supplemented with Kanamycine (50mg.L^−1^) and Chloramphenicol (34mg.L^−1^). Cultures were first incubated at 37°C to a 0.6 optical density at 600nm wavelength, then transferred and incubated at 16°C. When optical density reached 0.8, protein expression was induced using IPTG (0.5mM).

After overnight incubation at 16°C, cells were harvested at 6000 rpm for 10 minutes, washed once with cold 1x PBS, then collected at 5000 rpm for 10 minutes.

Cells were mixed in a lysis buffer (1.5x PBS, 20mM Imidazole, 0.1% Igepal, 10% Glycerol, 1mM MgCl2) supplemented with protease inhibitors (Aprotinin 2μg.mL^−1^, Leupeptin 1μg.mL^−1^, Pepstatin 1μg.mL^−1^, PMSF 50μg.mL^−1^) then lysed using sonication for 4 minutes on ice, and centrifuged at 18000 rpm for 25 minutes at 4°C.

Supernatants were mixed with 500μl Ni-NTA agarose resin (=500μl of 50% slurry, Clontech) pre-equilibrated in lysis buffer, and incubated for 2 hours in 50ml falcons on a rotator at 4 °C. Beads were collected and washed with lysis buffer then transferred on Bio-Spin columns (Biorad) pre-washed in lysis buffer. Columns were further washed with lysis buffer followed by a wash buffer (1.5x PBS, 50mM Imidazole, 250mM NaCl, 0.1% Igepal, 10% Glycerol, 1mM MgCl_2_), before protein elution in 800 ul fractions (1.5x PBS, 150mM Imidazole, 10% Glycerol, 1mM MgCl_2_).

Nmd4 was further dialyzed against calmodulin binding buffer (1x PBS, 100mM NaCl, 0.1% Igepal, 10% Glycerol, 1mM MgCl2, 4mM CaCl2, 1mM DTT) overnight at 4°C in Spectrapor-4 (12-14 MWCO), then mixed with 500μl Calmodulin Affinity Resin (= 1ml of 50% slurry, Agilent) pre-equilibrated in binding buffer. After 1 hour incubation into a Bio-Spin column on a rotator at 4°C, beads were washed twice with binding buffer, before protein elution (1x PBS, 100mM NaCl, 0.05% Igepal, 10% Glycerol, 1mM MgCl_2_, 20mM EGTA, 1mM DTT).

Proteins were finally dialyzed against 1.5x PBS, 150mM NaCl, 10% (w/v) glycerol, 1mM DTT and 1mM MgCl2 in Spectrapor-4 (12-14 MWCO) then stored at-80°C.

### *In vitro* pull-down assays

Pull-down was performed using preblocked calmodulin affinity beads (Agilent).

**Preblocking beads**. Briefly, in order to preblock beads, 1 ml calmodulin sepharose beads (50% Slurry) were spun, and resuspended in 20mM Hepes, 500mM NaCl, 0.1% Igepal, 0.08mg.ml^−1^ glycogen carrier, 0.08mg.ml^−1^ tRNA and 0.8mg.ml^−1^ BSA. After 2 hours at 4°C, beads were washed 3 times (20mM Hepes, 150mM NaCl, 0.1% Igepal) then resuspended in 500μl 1x binding buffer 250/10 (20mM Hepes, 250mM NaCl, 0.1% Igepal, 2mM MgAc_2_, 2mM CaCl_2_, 10% glycerol, 1mM DTT).

***In vitro* pull-down assay**. Recombinant CBP-Nmd4, yeast Upf1 and human Upf1 proteins were thawed on ice. Each mix contained 2μg of each protein in 30μl reaction mixes. NaCl and glycerol concentrations were adjusted to 150mM and 15% respectively. Five microliters (1/6) aliquots were taken out of each mix to load on Input gel. Each mix was further supplemented with 5μl of water, and 30μl of a NaCl/glycerol buffer to reach a 60μl reaction volume containing 125mM NaCl and 12.5 % glycerol.

Mixes were incubated for 20 minutes at 30°C. To perform pull-downs, 12μl of preblocked calmodulin beads were added to each mix, along with 200μl 1x binding buffer 150/10 (20mM Hepes, 150mM NaCl, 0.1% Igepal, 2mM MgAc_2_, 2mM CaCl_2_, 10% glycerol, 1mM DTT), then rotated 1 hour at 4°C. Beads were further washed 3 times with 1x binding buffer 150/10, then dried using a thinned Pasteur pipet. The retained complexes were eluted using 20μl elution buffer (10mM Tris pH=7.5, 150mM NaCl, 1mM MgAc_2_, 2mM Imidazole, 20mM EGTA, 0.1% Igepal, 14% glycerol, 10mM β-mercaptoethanol) while shaking 5 minutes at 1400 rpm, 30°C.

Eluates were collected after spinning, transferred to fresh tubes, concentrated 30 minutes in a Speed-vac then loaded on 10% SDS-PAGE gels.

### RNA extraction

For RT-qPCR and RNAseq, cells were first grown in YPD to log phase and collected. Total RNA was extracted using the hot phenol extraction method and precipitated using ammonium acetate and ethanol. The extracted RNA samples were treated with DNase I (Ambion TURBO DNA-free kit) before reverse-transcription (RT) and library preparation.

### Reverse transcription and quantitative PCR

To measure the enrichment of NMD substrates such as pre-messenger RPL28 or DAL7 in different experiments, we performed RT-qPCR. After RNA extraction and DNase treatment, 500ng of RNA were reverse-transcribed using Superscript III (Invitrogen) and a mix of reverse qPCR oligonucleotides, CS888 (RPL28-premature), CS889 (RPL28), CS1077 (RIM1), CS1430 (DAL7); for RNA immuno-precipitation we recovered 300ng to 3000ng and then use everything for reverse-transcription. cDNA were diluted to 1/8, 1/64, 1/512 and 1/4096 and 5μL were mixed 5μL with Eurobio green One step loRox mix and qPCR primers in a final volume of 25μL. The amplification was done in stratagene MX3005P and corresponding software (MXpro quantitative PCR system), according to the following program: cycle 1 (95°C for 5min) and cycle 2 (40 cycles of 95°C for 20s and 60°C for 1min). We chose the threshold manually and analyzed the results with home made excel sheet according the ΔΔCt method. Using the dilution curve, we calculated ΔCt that are the difference of Ct between the RNA tested and the reference RNA; e.g. ΔCt = Ct mRNA of interest / Ct reference RNA. We next calculated the ΔΔCt to normalize the values to the WT condition; for the comparison of NMD substrates expression in total RNA extract, we compared the ΔCt of the mutant with the ΔCt of the WT and we calculated the NMD efficiency based on these ΔΔCt; for RNA immuno-precipitated the ΔΔCt were the difference between the Ct of immuno-precipitation and input.

### NMD efficiency calculation

Complementation of an NMD deficient strain leads to a decrease in the levels of an NMD-sensitive transcript like RPL28. To be able to judge how the efficiency of NMD affects the level of a transcript and to simplify the interpretation of complementation experiments, we developed a formula to calculate NMD efficiency using the maximum and minimum steady-state levels of a given transcript. We assume that RNA synthesis is constant and that the degradation of RNA occurs by two competitive pathways, with different first-order rate constants: k_NMD_ and k_base_. Thus, the variation in the RNA levels with time could be expressed as:

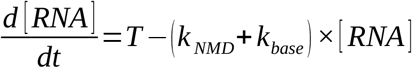

where:

T is transcription rate (synthesis, and export),
[RNA] is RNA concentration,
k_NMD_ is the global rate constant for degradation through NMD,
k_base_ is the base rate constant for degradation of the RNA, independent of NMD.

At steady state, the change in RNA concentration is null. A given steady-state level can be obtained either by both high transcription and high degradation or low transcription and low degradation. Steady state levels of a given RNA would be:

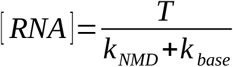

Since most of the time we do not know the transcription rate, or the degradation constants, we need to remove one of the variables. Let us consider two situations in which transcription is supposed invariable, a wt and a mutant strain devoid of NMD:

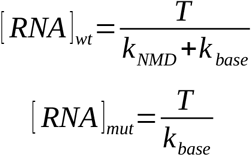

Thus, the ratio between the mutant that has no NMD and wild type, N, is:

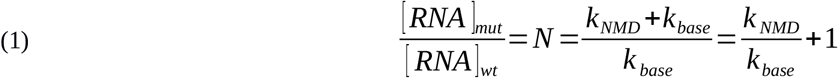

If the RNA is not an NMD substrate, kNMD is 0 and the ratio between the mutant and wt becomes 1.

If the RNA is exclusively degraded through NMD, the ratio would be infinite.

At a fractional NMD efficiency, noted α, the degradation through NMD would be α × kNMD and the ratio of RNA towards wt in this mutant, P, becomes:

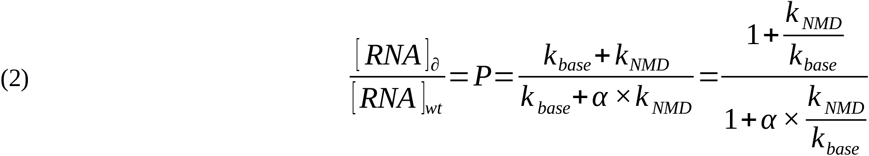

N is always larger than P.

We can substitute *k*_*NMD*_/*k*_*base*_in (2) with *N*−1, from (1):

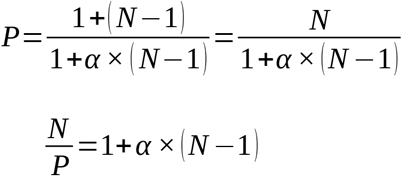

Thus the fraction of NMD, used to define efficiency, can be calculated from:

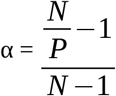

Where N is the enrichment of RNA in the NMD null mutant and P the enrichment in the tested strain.

Example: if the RNA is enriched 8 fold in an NMD mutant and 4 fold in a “partial” NMD strain over WT, the fractional efficiency of NMD would be:

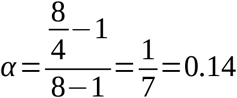

This example shows that the correlation between observed ratios and NMD efficiency is not linear. The formula maps values obtained by RT-QPCR to a percentage of “NMD efficiency” and works fine for extreme values. If a mutant has no NMD defect, P will be equal to 1 and α becomes 1. If the mutant is 100% NMD deficient, P will approach N and, α becomes 0. Due to experimental error, P might be superior to N, but in that case α should be considered 0.

### RNA libraries preparation, sequencing

RNA, extracted and treated with DNase, were subjected to Ribozero^®^ treatment to remove ribosomal RNAs. Removal was verified by testing the RNAs on a Bioanalyser. Libraries of mRNA were prepared with a TruSeq Stranded kit (Illumina). We did the protocol from the fragmentation step to the amplification. For *nmd4*Δ, *ebs1*Δ, *nmd4*Δ/*ebs1*Δ and the BY4741 associated, the fragmentation incubation was reduced to 7 minutes, and we amplified the library with 10 cycles only. Libraries were sequenced on the Illumina HiSeq 2500 for 50 bases (for upf1Δ and BY4741 associated) and 65 bases (for *nmd4*Δ, *ebs1*Δ, *nmd4*Δ/*ebs1*Δ and BY4741 associated). Sequenced reads were aligned to yeast genome version sacCer3 (version R64-2-1) using STAR (Dobin *et al*, 2013) with the default parameters except for-s 0-0. We used feature_count from the subread package (version 1.5.0-p3-Linux-x86_64) to count the number of reads per features. For the features, we used a custom list where cryptic transcripts introns could be specifically counted. There were three independent experiments for each condition; *upf1*Δ and the first BY4741 and *nmd4*Δ, *ebs1*Δ, *nmd4*Δ/*ebs1*Δ and the second BY4741 were done by different experimenters and were not sequenced in the same run.

### Polysome fractionation

Polysome extracts were obtained from 150mL of mid-log phase (*A*_*600*_ ≈ 0.6) yeast cells culture treated with 50μg·ml^−1^ cycloheximide for 5 minutes at 4°C and broken with glass beads using a MagNA lyser (three times for 60s at 3000 rpm) in breaking buffer (20mM Tris-HCl, pH 7.4, 50mM KCl, 10mM MgCl2, 50μg·ml^−1^ cycloheximide, 1mM DTT, 0.1mg·ml ^−1^ Heparin, and EDTA-free protease inhibitors from Roche). 10 to 15 *A*260 of clarified cell lysates were loaded on 10–50% sucrose gradients. Gradients were centrifuged 3h15 at 36000 rpm with slow acceleration in a SW41-Ti rotor (Beckman) in a Optima L-80 ultracentrifuge (Beckman). Half of collected fractions were precipitated with TCA and the precipitate was resuspended in 50μL of sample buffer. 10μL were loaded on polyacrylamide NuPAGE Novex 4-12% Bis-Tris gels (Life technologies). After transfer to nitrocellulose membrane with a semi-dry system, proteins were detected by hybridization with appropriate antibodies.

### Protein sequence alignment, phylogenetic trees and percentage of identity calculation

We used metaPhOr (Pryszcz *et al*, 2011) to obtain sequences of orthologues for the studied proteins. We aligned entire proteins or fragments using Mafft software called as a web service from Jalview. PIN domain boundaries for Smg6, Smg5 have been determined by sequence homology with orthologous proteins from *D. melanogaster*, *C. elegans*, *M. musculus* (positions 1239-1419 for hSmg6, 849-1016 for hSmg5). For Nmd4, we used all the 1-218 amino acid sequence for the alignment, even if the last 60 amino acids are not part of the PIN domain. Identity was computed as the percentage of identical aligned residues over the total number of aligned residues. To strengthen alignment reliability, we also performed delta-BLAST alignment (Boratyn *et al*, 2012) on the *S. cerevisiae* protein database with the hSmg6 and hSmg5 PIN domains and hSmg7 14-3-3 domain as query, using E-values as significance scores.

## QUANTIFICATION AND STATISTICAL ANALYSIS

### Western blot quantification

For polysome analysis, western blot signals were quantified using Image Lab software (version 5.2.1 build 11 – Biorad). We used the square volume tool to define boxes of identical size at the expected position of the protein band we wanted to quantify. We also defined a series of blank boxes for each lane for background definition. To normalize the results, we calculated the percentage of signal over all the fractions of a run for both Nmd4 and Rps8 (control). Statistical significance of the signal differences between conditions were assessed by the paired Student t-test. Stars on plots indicate that the p-value associated with the test was less than 0.05.

### GO Term analysis and statistics

We used the GO Term Finder tool associated with SGD database (Cherry *et al*, 2012) to search for common function of protein list extracted from MS analysis; for example, the group of proteins removed by RNase treatment and group of proteins of a given enrichment; default settings were used. We calculated the significance of the accumulation of certain protein classes using hypergeometric distribution test (**Supplementary Table 5**). This test allows determining if the sampling is random or if there is an enrichment of specific classes of annotations.

### Mass spectrometry results analysis

Raw data files from MS were searched using the MaxQuant/Andromeda search engine (version 1.5.5.1 (Cox & Mann, 2008; Cox *et al*, 2011) against the Uniprot *Saccharomyces cerevisiae* proteome database (downloaded 12/02/2016; total 6,721 entries) completed with the 22 virus protein sequences from yeast, and a list of commonly observed contaminants supplied by MaxQuant. Searches allowed either trypsin specificity with two missed cleavages. Cysteine carbamidomethylation was selected as fixed modification, and oxidation of methionine and N-terminal acetylation were searched as variable modifications. Peptide identification was performed using 6 ppm as mass tolerance at the MS level for FT mass analyser and 0.5Da at the MS/MS level for IT analyzer. MaxQuant/Andromeda used seven amino acid minimum for peptide length, with 0.01 false discovery rate for both protein and peptide identification. The protein identification was considered valid only when at least one unique or “razor” peptide was present. Following protein identification, the intensity for each identified protein was calculated using peptide signal intensities. Identification transfer protocol (“match-between-runs” feature in MaxQuant) was applied.

Only protein identifications based on a minimum of two peptides were selected for further quantitative studies. Bioinformatics analysis of the MaxQuant/Andromeda workflow output and the analysis of the abundances of the identified proteins were performed with custom R scripts (**Supplementary Fig. 1**). R scripts were used to calculate a score called LTOP2 for each protein group in the different experiments. This score is similar with the “top three” average described (Silva *et al*, 2006), with several differences. First, we built meta-peptide intensities, based on the intensity of overlapping peptides with missed cleavages (minimum 4 residues overlapping). Next, we took the 3 most intense peptides (top3) or the top2 if only two peptide intensities were available, and calculated the average of log2-transformed intensities. LTOP2 values on whole cell extracts were in excellent agreement with previous estimates of protein abundance in yeast (**Fig. 1d**). Finally, LTOP2 between experiments were normalized using the TEV protease as a spike in, as this protease is added with the same amount in each sample during the purification process. This procedure allows us to compare all the experiments. To overcome some limitations of the score, illustrated in Fig. 1e, we calculated enrichments for each protein group. These enrichments were the ratio of the proportion of each protein in the purification against the proportion in the input. Using LTOP2, we calculated relative levels of each protein in the samples, noted L. With Pp the proportion of a protein group in purification, Pi the proportion of the protein group in the input and X the protein group, the enrichment (E) will be calculated as follows:

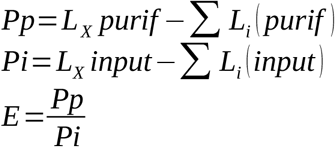

Enrichments were computed for each protein in each experiment, transformed to log2 values (**Supplementary Table 3**) and served as base to calculate enrichment mean and standard deviation (**Supplementary Table 4**) that we used for graphical representations.

### RNAseq statistical analysis

Expression of gene in the different samples and replicates were normalized to WT strain using DESeq2 (Love *et al*, 2014). We analyzed the three replicates independently and used the mean for figures. During DESeq2 workflow, we removed features identified by zero reads to avoid problems with the logarithm transformation. We determined NMD substrates from *upf1*Δ RNAseq results and fixed the threshold for these RNA to a minimum of 1.4 fold increase. For the bin analysis, we used a custom script to divide RNA sequenced in *nmd4*Δ and *ebs1*Δ experiments into 5 bins containing the same number of transcript and calculate the percentage of NMD substrates for each bin ( **Fig. 5**). Statistical significance of the differences between bins was assessed using a binomial test.

## DATA AND SOFTWARE AVAILABILITY

In addition to the access to raw data, enrichment and intensity data are available as an interactive web application at: https://hub05.hosting.pasteur.fr/NMD_complexes.

## Strains used in this study

Dehecq et al., Detection and degradation of nonsense-mediated mRNA decay substrates involve two distinct Upf1-bound complexes

| strain | original strain | Genotype | Reference |
| --- | --- | --- | --- |
| LMA2154 (BY4741) |  | *MATa ura3Δ0 his3Δ1 leu2Δ0 met15Δ0* | Brachmann et al. 1998 |
| LMA2155 (BY4742) |  | *MATa ura3Δ0 his3Δ1 leu2Δ0 lys2Δ0* | Brachmann et al. 1998 |
| LMA2194 (UPF1-TAP) | BY4741 | NAM7-TAP::HIS3MX | Ghaemmighami et al., 2003 |
| LMA3730 (UPF1-TAP) | BY4741 | NAM7-CRAP::URA3 | This study |
| LMA2192 (UPF2-TAP) | BY4741 | NMD2-TAP::HIS3MX | Ghaemmighami et al., 2003 |
| LMA3731 (UPF2-TAP) | BY4741 | NMD2-CRAP::URA3 | This study |
| LMA2193 (UPF3-TAP) | BY4741 | UPF3-TAP::HIS3MX | Ghaemmighami et al., 2003 |
| LMA4263 (UPF3-TAP) | BY4741 | UPF3-CRAP::URA3 | This study |
| LMA3317 (NMD4-TAP) | BY4741 | NMD4-TAP::HIS3MX | Ghaemmighami et al., 2003 |
| LMA3728/3729 (NMD4-TAP) | BY4741 | NMD4-CRAP::URA3 | This study |
| LMA4264 (DCP1-TAP) | BY4741 | DCP1-TAP::HIS3MX | Ghaemmighami et al., 2003 |
| (EBS1-TAP) | BY4741 | EBS1-CRAP::URA3 | This study |
| LMA3312 (HRR25-TAP) | BY4741 | HRR25-TAP::HIS3MX | Ghaemmighami et al., 2003 |
| LMA3849 (UPF1-TAP/NMD4-HA) | BY4741 | NAM7-CRAP::URA3 / NMD4-HA::KANMX6 | This study |
| LMA3851 (UPF1-TAP/EBS1-HA) | BY4741 | NAM7-CRAP::URA3 / EBS1-HA::KANMX6 | This study |
| LMA3852 (UPF1-TAP/EDC3) | BY4741 | NAM7-CRAP::URA3 / EDC3-HA::KANMX6 | This study |
| LMA1667 (upf1Δ) | BY4741 | NAM7::KANMX6 | Giaever et al., 2002 |
| LMA1669 (upf2Δ) | BY4741 | NMD2::KANMX6 | Giaever et al., 2002 |
| LMA1671 (upf3Δ) | BY4741 | UPF3::KANMX6 | Giaever et al., 2002 |
| LMA3732 (nmd4Δ) | BY4741 | NMD4::KANMX6 | Giaever et al., 2002 |
| LMA4112 (UPF2-TAP/upf1Δ) | BY4741 | NAM7::KANMX6 / NMD2-CRAP::URA3 | This study |
| LMA4113 (UPF3-TAP/upf1Δ) | BY4741 | NAM7::KANMX6 / UPF3-CRAP::URA3 | This study |
| LMA4114 (NMD4-TAP/upf1Δ) | BY4741 | NAM7::KANMX6 / NMD4-CRAP::URA3 | This study |
| LMA3739 (UPF1-TAP/upf2Δ) | BY4741 | NMD2::KANMX6 / NAM7-CRAP::URA3 | This study |
| LMA4701/4702 (NMD4-TAP/upf2Δ) | BY4741 | NMD2::KANMX6 / NMD4-CRAP::URA3 | This study |
| LMA3735 (UPF1-TAP/upf3Δ) | BY4741 | UPF3::KANMX6 / NAM7-CRAP::URA3 | This study |
| LMA3736 (UPF1-TAP/nmd4Δ) | BY4741 | NMD4::KANMX6 / NAM7-CRAP::URA3 | This study |
| LMA4523 (upf1Δ/upf2Δ) | BY4741 | NAM7::HIS3MX / NMD2::KANMX6 | This study |
| LMA4524 (upf1Δ/upf3Δ) | BY4741 | NAM7::HIS3MX / UPF3::KANMX6 | This study |
| LMA4525 (upf1Δ/ebs1Δ) | BY4741 | NAM7::HIS3MX / EBS1::KANMX6 | This study |
| LMA4678 (upf1Δ/nmd4Δ) | BY4741 | NAM7::HIS3MX / NMD4::KANMX6 | This study |
| LMA3853/3854 (nmd4Δ/ebs1Δ) | BY4742 | NMD4::ProMFalpha2NAT / EBS1::KANMX6 | This study |

## Plasmids used in this study

| Plasmid | Description and ID (yeast marker) | Reference |
| --- | --- | --- |
| pRS316 | pRS316 (URA3) | Sikorski & Hieter, 1989 |
| pCM189-NTAP | pl.1233 (H1) (URA3) | This study |
| pCM189-NTAP-NAM7-FL | pl.1442 (TAP-UPF1-FL) (URA3) | This study |
| pCM189-NTAP-NAM7-CH | pl.1443 (TAP-UPF1-CH) (URA3) | This study |
| pCM189-NTAP-NAM7-NoCH | pl.1444 (TAP-UPF1-Cter) (URA3) | This study |
| pCM189-NTAP-Upf1 2-853 | pl.1521 (TAP-UPF1-NoSQ) (URA3) | This study |
| pCM189-NTAP-Upf1 208-853 | pl.1522 (TAP-UPF1-Cter-NoSQ) (URA3) | This study |
| pDEST14-NAM7 | pl.1350 – Gateway destination vector | This study |
| pDONR201-NAM7 | pl.1330 – Gateway source vector | This study |
| pCRBlunt-CRAP(6-HisTAP) | pl.1287 – TAP to CRAP cassette vector | This study |

## Oligonucleotides used in this study

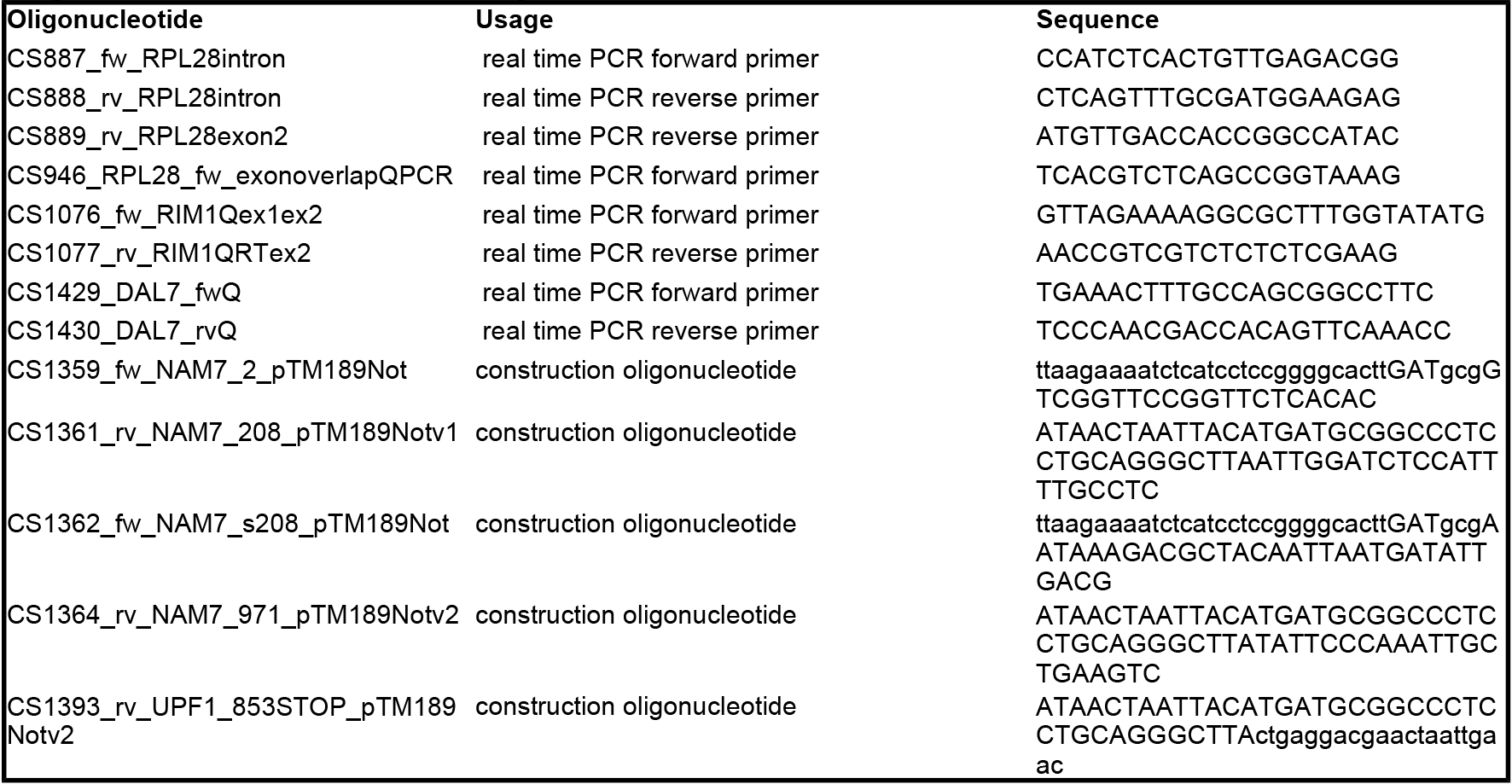

## References

Addinall SG, Holstein E-M, Lawless C, Yu M, Chapman K, Banks AP, Ngo H-P, Maringele L, Taschuk M, Young A, Ciesiolka A, Lister AL, Wipat A, Wilkinson DJ & Lydall D (2011) Quantitative fitness analysis shows that NMD proteins and many other protein complexes suppress or enhance distinct telomere cap defects. PLoS Genet. 7: e1001362

Ahrné E, Molzahn L, Glatter T & Schmidt A (2013) Critical assessment of proteome-wide label-free absolute abundance estimation strategies. Proteomics 13: 2567–2578

Amrani N, Ganesan R, Kervestin S, Mangus DA, Ghosh S & Jacobson A (2004) A faux 3’-UTR promotes aberrant termination and triggers nonsense-mediated mRNA decay. Nature 432: 112–118

Atkin AL, Schenkman LR, Eastham M, Dahlseid JN, Lelivelt MJ & Culbertson MR (1997) Relationship between yeast polyribosomes and Upf proteins required for nonsense mRNA decay. J. Biol. Chem. 272: 22163–22172

Azzalin CM, Reichenbach P, Khoriauli L, Giulotto E & Lingner J (2007) Telomeric repeat containing RNA and RNA surveillance factors at mammalian chromosome ends. Science 318: 798–801

Behm-Ansmant I, Gatfield D, Rehwinkel J, Hilgers V & Izaurralde E (2007) A conserved role for cytoplasmic poly(A)-binding protein 1 (PABPC1) in nonsense-mediated mRNA decay. EMBO J. 26: 1591–1601

Benschop JJ, Brabers N, van Leenen D, Bakker LV, van Deutekom HWM, van Berkum NL, Apweiler E, Lijnzaad P, Holstege FCP & Kemmeren P (2010) A Consensus of Core Protein Complex Compositions for Saccharomyces cerevisiae. Molecular Cell 38: 916–928

Brannan KW, Jin W, Huelga SC, Banks CAS, Gilmore JM, Florens L, Washburn MP, Van Nostrand EL, Pratt GA, Schwinn MK, Daniels DL & Yeo GW (2016) SONAR Discovers RNA-Binding Proteins from Analysis of Large-Scale Protein-Protein Interactomes. Molecular Cell 64: 282–293

Bühler M, Steiner S, Mohn F, Paillusson A & Mühlemann O (2006) EJC-independent degradation of nonsense immunoglobulin-μ mRNA depends on 3′ UTR length. Nat Struct Mol Biol 13: 462–464

Chakrabarti S, Bonneau F, Schüssler S, Eppinger E & Conti E (2014) Phospho-dependent and phospho-independent interactions of the helicase UPF1 with the NMD factors SMG5–SMG7 and SMG6. Nucl. Acids Res.: gku578

Chamieh H, Ballut L, Bonneau F & Le Hir H (2008) NMD factors UPF2 and UPF3 bridge UPF1 to the exon junction complex and stimulate its RNA helicase activity. Nat Struct Mol Biol 15: 85–93

Chapin A, Hu H, Rynearson SG, Hollien J, Yandell M & Metzstein MM (2014) In Vivo Determination of Direct Targets of the Nonsense-Mediated Decay Pathway in Drosophila. G3 (Bethesda) 4: 485–496

Chen Z, Smith KR, Batterham P & Robin C (2005) Smg1 Nonsense Mutations Do Not Abolish Nonsense-Mediated mRNA Decay in Drosophila melanogaster. Genetics 171: 403–406

Clerici M, Mourao A, Gutsche I, Gehring NH, Hentze MW, Kulozik A, Kadlec J, Sattler M & Cusack S (2009) Unusual bipartite mode of interaction between the nonsense-mediated decay factors, UPF1 and UPF2. EMBO J 28: 2293–2306

Clissold PM & Ponting CP (2000) PIN domains in nonsense-mediated mRNA decay and RNAi. Current Biology 10: R888–R890

Collins SR, Kemmeren P, Zhao X-C, Greenblatt JF, Spencer F, Holstege FCP, Weissman JS & Krogan NJ (2007) Toward a Comprehensive Atlas of the Physical Interactome of Saccharomyces cerevisiae. Mol Cell Proteomics 6: 439–450

Colombo M, Karousis ED, Bourquin J, Bruggmann R & Mühlemann O (2016) Transcriptome-wide identification of NMD-targeted human mRNAs reveals extensive redundancy between SMG6-and SMG7-mediated degradation pathways. RNA: rna.059055.116

Cui Y, Hagan KW, Zhang S & Peltz SW (1995) Identification and characterization of genes that are required for the accelerated degradation of mRNAs containing a premature translational termination codon. Genes Dev. 9: 423–436

Dujon B, Sherman D, Fischer G, Durrens P, Casaregola S, Lafontaine I, De Montigny J, Marck C, Neuvéglise C, Talla E, Goffard N, Frangeul L, Aigle M, Anthouard V, Babour A, Barbe V, Barnay S, Blanchin S, Beckerich J-M, Beyne E, et al (2004) Genome evolution in yeasts. Nature 430: 35–44

Eberle AB, Lykke-Andersen S, Mühlemann O & Jensen TH (2009) SMG6 promotes endonucleolytic cleavage of nonsense mRNA in human cells. Nat Struct Mol Biol 16: 49–55

Eberle AB, Stalder L, Mathys H, Orozco RZ & Mühlemann O (2008) Posttranscriptional gene regulation by spatial rearrangement of the 3’ untranslated region. PLoS Biol. 6: Available at: http://www.ncbi.nlm.nih.gov/pubmed/18447580 [Accessed September 18, 2012]

Gatfield D, Unterholzner L, Ciccarelli FD, Bork P & Izaurralde E (2003) Nonsense-mediated mRNA decay in Drosophila:at the intersection of the yeast and mammalian pathways. EMBO J 22: 3960–3970

Gavin A-C, Aloy P, Grandi P, Krause R, Boesche M, Marzioch M, Rau C, Jensen LJ, Bastuck S, Dümpelfeld B, Edelmann A, Heurtier M-A, Hoffman V, Hoefert C, Klein K, Hudak M, Michon A-M, Schelder M, Schirle M, Remor M, et al (2006) Proteome survey reveals modularity of the yeast cell machinery. Nature 440: 631–636

Ghosh S, Ganesan R, Amrani N & Jacobson A (2010) Translational competence of ribosomes released from a premature termination codon is modulated by NMD factors. RNA 16: 1832–1847

Halbeisen RE, Scherrer T & Gerber AP (2009) Affinity purification of ribosomes to access the translatome. Methods 48: 306–310

Hall GW & Thein S (1994) Nonsense codon mutations in the terminal exon of the beta-globin gene are not associated with a reduction in beta-mRNA accumulation: a mechanism for the phenotype of dominant beta-thalassemia. Blood 83: 2031–2037

He F, Brown AH & Jacobson A (1996) Interaction between Nmd2p and Upf1p is required for activity but not for dominant-negative inhibition of the nonsense-mediated mRNA decay pathway in yeast. RNA 2: 153–170

He F, Brown AH & Jacobson A (1997) Upf1p, Nmd2p, and Upf3p are interacting components of the yeast nonsense-mediated mRNA decay pathway. Mol Cell Biol 17: 1580–1594

He F & Jacobson A (1995) Identification of a novel component of the nonsense-mediated mRNA decay pathway by use of an interacting protein screen. Genes Dev. 9: 437–454

He F & Jacobson A (2001) Upf1p, Nmd2p, and Upf3p regulate the decapping and exonucleolytic degradation of both nonsense-containing mRNAs and wild-type mRNAs. Mol. Cell. Biol. 21: 1515–1530

He F & Jacobson A (2015) Nonsense-Mediated mRNA Decay: Degradation of Defective Transcripts Is Only Part of the Story. Annu. Rev. Genet. 49: 339–366

Heyer EE & Moore MJ (2016) Redefining the Translational Status of 80S Monosomes. Cell 164: 757–769

Ho B, Baryshnikova A & Brown GW (2018) Unification of Protein Abundance Datasets Yields a Quantitative Saccharomyces cerevisiae Proteome. Cell Systems Available at: http://www.sciencedirect.com/science/article/pii/S240547121730546X [Accessed February 5, 2018]

Holbrook JA, Neu-Yilik G, Hentze MW & Kulozik AE (2004) Nonsense-mediated decay approaches the clinic. Nat Genet 36: 801–808

Huang L, Lou C-H, Chan W, Shum EY, Shao A, Stone E, Karam R, Song H-W & Wilkinson MF (2011) RNA Homeostasis Governed by Cell Type-Specific and Branched Feedback Loops Acting on NMD. Mol Cell 43: 950–961

Hug N & Cáceres JF (2014) The RNA Helicase DHX34 Activates NMD by Promoting a Transition from the Surveillance to the Decay-Inducing Complex. Cell Rep 8: 1845–1856

Huntzinger E, Kashima I, Fauser M, Sauliere J & Izaurralde E (2008) SMG6 is the catalytic endonuclease that cleaves mRNAs containing nonsense codons in metazoan. RNA 14: 2609–2617

Johansson MJO, He F, Spatrick P, Li C & Jacobson A (2007) Association of yeast Upf1p with direct substrates of the NMD pathway. Proc Natl Acad Sci U S A 104: 20872–20877

Johns L, Grimson A, Kuchma SL, Newman CL & Anderson P (2007) Caenorhabditis elegans SMG-2 Selectively Marks mRNAs Containing Premature Translation Termination Codons. Mol Cell Biol 27: 5630–5638

Kashima I, Yamashita A, Izumi N, Kataoka N, Morishita R, Hoshino S, Ohno M, Dreyfuss G & Ohno S (2006) Binding of a novel SMG-1–Upf1–eRF1–eRF3 complex (SURF) to the exon junction complex triggers Upf1 phosphorylation and nonsense-mediated mRNA decay. Genes Dev. 20: 355–367

Krogan NJ, Cagney G, Yu H, Zhong G, Guo X, Ignatchenko A, Li J, Pu S, Datta N, Tikuisis AP, Punna T, Peregrín-Alvarez JM, Shales M, Zhang X, Davey M, Robinson MD, Paccanaro A, Bray JE, Sheung A, Beattie B, et al (2006) Global landscape of protein complexes in the yeast Saccharomyces cerevisiae. Nature 440: 637–643

Lasalde C, Rivera AV, León AJ, González-Feliciano JA, Estrella LA, Rodríguez-Cruz EN, Correa ME, Cajigas IJ, Bracho DP, Vega IE, Wilkinson MF & González CI (2013) Identification and functional analysis of novel phosphorylation sites in the RNA surveillance protein Upf1. Nucleic Acids Res.

Le Hir H, Gatfield D, Izaurralde E & Moore MJ (2001) The exon–exon junction complex provides a binding platform for factors involved in mRNA export and nonsense-mediated mRNA decay. EMBO J 20: 4987–4997

Leeds P, Peltz SW, Jacobson A & Culbertson MR (1991) The product of the yeast UPF1 gene is required for rapid turnover of mRNAs containing a premature translational termination codon. Genes Dev. 5: 2303–2314

Leeds P, Wood JM, Lee BS & Culbertson MR (1992) Gene products that promote mRNA turnover in Saccharomyces cerevisiae. Mol. Cell. Biol. 12: 2165–2177

Lindeboom RGH, Supek F & Lehner B (2016) The rules and impact of nonsense-mediated mRNA decay in human cancers. Nat Genet 48: 1112–1118

Lloyd JPB & Davies B (2013) SMG1 is an ancient nonsense-mediated mRNA decay effector. Plant J 76: 800–810

Loh B, Jonas S & Izaurralde E (2013) The SMG5–SMG7 heterodimer directly recruits the CCR4– NOT deadenylase complex to mRNAs containing nonsense codons via interaction with POP2. Genes Dev 27: 2125–2138

Love MI, Huber W & Anders S (2014) Moderated estimation of fold change and dispersion for RNA-seq data with DESeq2. Genome Biology 15: 550

Luke B, Azzalin CM, Hug N, Deplazes A, Peter M & Lingner J (2007) Saccharomyces cerevisiae Ebs1p is a putative ortholog of human Smg7 and promotes nonsense-mediated mRNA decay. Nucleic Acids Res. 35: 7688–7697

Lykke-Andersen J (2002) Identification of a Human Decapping Complex Associated with hUpf Proteins in Nonsense-Mediated Decay. Mol Cell Biol 22: 8114–8121

Lykke-Andersen J, Shu M-D & Steitz JA (2001) Communication of the Position of Exon-Exon Junctions to the mRNA Surveillance Machinery by the Protein RNPS1. Science 293: 1836–1839

Lykke-Andersen S, Chen Y, Ardal BR, Lilje B, Waage J, Sandelin A & Jensen TH (2014) Human nonsense-mediated RNA decay initiates widely by endonucleolysis and targets snoRNA host genes. Genes Dev 28: 2498–2517

Malabat C, Feuerbach F, Ma L, Saveanu C & Jacquier A (2015) Quality control of transcription start site selection by nonsense-mediated-mRNA decay. Elife 4:

Maquat LE (2004) Nonsense-mediated mRNA decay: splicing, translation and mRNP dynamics. Nat. Rev. Mol. Cell Biol. 5: 89–99

Medghalchi SM, Frischmeyer PA, Mendell JT, Kelly AG, Lawler AM & Dietz HC (2001) Rent1, a trans-effector of nonsense-mediated mRNA decay, is essential for mammalian embryonic viability. Hum. Mol. Genet. 10: 99–105

Mellacheruvu D, Wright Z, Couzens AL, Lambert J-P, St-Denis NA, Li T, Miteva YV, Hauri S, Sardiu ME, Low TY, Halim VA, Bagshaw RD, Hubner NC, al-Hakim A, Bouchard A, Faubert D, Fermin D, Dunham WH, Goudreault M, Lin Z-Y, et al (2013) The CRAPome: a contaminant repository for affinity purification-mass spectrometry data. Nat Meth 10: 730–736

Metze S, Herzog VA, Ruepp M-D & Mühlemann O (2013) Comparison of EJC-enhanced and EJC-independent NMD in human cells reveals two partially redundant degradation pathways. RNA 19: 1432–1448

Muhlrad D & Parker R (1994) Premature translational termination triggers mRNA decapping. Nature 370: 578–581

Muhlrad D & Parker R (1999) Aberrant mRNAs with extended 3’ UTRs are substrates for rapid degradation by mRNA surveillance. RNA 5: 1299–1307

Nelson JO, Moore KA, Chapin A, Hollien J & Metzstein MM (2016) Degradation of Gadd45 mRNA by nonsense-mediated decay is essential for viability. eLife 5: Available at: https://www.ncbi.nlm.nih.gov/pmc/articles/PMC4848089/ [Accessed November 20, 2017]

Neu-Yilik G, Raimondeau E, Eliseev B, Yeramala L, Amthor B, Deniaud A, Huard K, Kerschgens K, Hentze MW, Schaffitzel C & Kulozik AE (2017) Dual function of UPF3B in early and late translation termination. The EMBO Journal: e201797079

Nicholson P, Josi C, Kurosawa H, Yamashita A & Mühlemann O (2014) A novel phosphorylation-independent interaction between SMG6 and UPF1 is essential for human NMD. Nucleic Acids Res 42: 9217–9235

Oeffinger M, Wei KE, Rogers R, DeGrasse JA, Chait BT, Aitchison JD & Rout MP (2007) Comprehensive analysis of diverse ribonucleoprotein complexes. Nat. Methods 4: 951–956

Ohnishi T, Yamashita A, Kashima I, Schell T, Anders KR, Grimson A, Hachiya T, Hentze MW, Anderson P & Ohno S (2003) Phosphorylation of hUPF1 Induces Formation of mRNA Surveillance Complexes Containing hSMG-5 and hSMG-7. Molecular Cell 12: 1187–1200

Okada-Katsuhata Y, Yamashita A, Kutsuzawa K, Izumi N, Hirahara F & Ohno S (2012) N-and C-terminal Upf1 phosphorylations create binding platforms for SMG-6 and SMG-5:SMG-7 during NMD. Nucleic Acids Res 40: 1251–1266

Page MF, Carr B, Anders KR, Grimson A & Anderson P (1999) SMG-2 Is a Phosphorylated Protein Required for mRNA Surveillance in Caenorhabditis elegans and Related to Upf1p of Yeast. Mol Cell Biol 19: 5943–5951

Perlick HA, Medghalchi SM, Spencer FA, Kendzior RJ & Dietz HC (1996) Mammalian orthologues of a yeast regulator of nonsense transcript stability. Proc Natl Acad Sci U S A 93: 10928–10932

Pryszcz LP, Huerta-Cepas J & Gabaldón T (2011) MetaPhOrs: orthology and paralogy predictions from multiple phylogenetic evidence using a consistency-based confidence score. Nucleic Acids Res 39: e32

Pulak R & Anderson P (1993) mRNA surveillance by the Caenorhabditis elegans smg genes. Genes Dev. 7: 1885–1897

Rigaut G, Shevchenko A, Rutz B, Wilm M, Mann M & Séraphin B (1999) A generic protein purification method for protein complex characterization and proteome exploration. Nat. Biotechnol 17: 1030–1032

Schell T, Köcher T, Wilm M, Seraphin B, Kulozik AE & Hentze MW (2003) Complexes between the nonsense-mediated mRNA decay pathway factor human upf1 (up-frameshift protein 1) and essential nonsense-mediated mRNA decay factors in HeLa cells. Biochem. J. 373: 775–783

Serdar LD, Whiteside DL & Baker KE (2016) ATP hydrolysis by UPF1 is required for efficient translation termination at premature stop codons. Nature Communications 7: 14021

Silva JC, Gorenstein MV, Li G-Z, Vissers JPC & Geromanos SJ (2006) Absolute Quantification of Proteins by LCMSE A Virtue of Parallel ms Acquisition. Mol Cell Proteomics 5: 144–156

Singh G, Rebbapragada I & Lykke-Andersen J (2008) A competition between stimulators and antagonists of Upf complex recruitment governs human nonsense-mediated mRNA decay. PLoS Biol. 6:

Swisher KD & Parker R (2011) Interactions between Upf1 and the Decapping Factors Edc3 and Pat1 in Saccharomyces cerevisiae. PLoS One 6: Available at: http://www.ncbi.nlm.nih.gov/pmc/articles/PMC3204985/ [Accessed July 12, 2017]

Tian M, Yang W, Zhang J, Dang H, Lu X, Fu C & Miao W (2017) Nonsense-mediated mRNA decay in Tetrahymena is EJC independent and requires a protozoa-specific nuclease. Nucleic Acids Res. 45: 6848–6863

Valásek L, Szamecz B, Hinnebusch AG & Nielsen KH (2007) In vivo stabilization of preinitiation complexes by formaldehyde cross-linking. Meth. Enzymol. 429: 163–183

Wen J & Brogna S (2010) Splicing-dependent NMD does not require the EJC in Schizosaccharomyces pombe. EMBO J 29: 1537–1551

Weng Y, Czaplinski K & Peltz SW (1996) Identification and characterization of mutations in the UPF1 gene that affect nonsense suppression and the formation of the Upf protein complex but not mRNA turnover. Mol Cell Biol 16: 5491–5506

Weng Y, Czaplinski K & Peltz SW (1998) ATP is a cofactor of the Upf1 protein that modulates its translation termination and RNA binding activities. RNA 4: 205–214

Yamashita A, Ohnishi T, Kashima I, Taya Y & Ohno S (2001) Human SMG-1, a novel phosphatidylinositol 3-kinase-related protein kinase, associates with components of the mRNA surveillance complex and is involved in the regulation of nonsense-mediated mRNA decay. Genes Dev 15: 2215–2228

Yepiskoposyan H, Aeschimann F, Nilsson D, Okoniewski M & Mühlemann O (2011) Autoregulation of the nonsense-mediated mRNA decay pathway in human cells. RNA 17: 2108–2118

Zhang S, Welch EM, Hogan K, Brown AH, Peltz SW & Jacobson A (1997) Polysome-associated mRNAs are substrates for the nonsense-mediated mRNA decay pathway in Saccharomyces cerevisiae. RNA 3: 234–244

## REFERENCES

Boratyn GM, Schäffer AA, Agarwala R, Altschul SF, Lipman DJ & Madden TL (2012) Domain enhanced lookup time accelerated BLAST. Biol Direct 7: 12

Cherry JM, Hong EL, Amundsen C, Balakrishnan R, Binkley G, Chan ET, Christie KR, Costanzo MC, Dwight SS, Engel SR, Fisk DG, Hirschman JE, Hitz BC, Karra K, Krieger CJ, Miyasato SR, Nash RS, Park J, Skrzypek MS, Simison M, et al (2012) Saccharomyces Genome Database: the genomics resource of budding yeast. Nucleic Acids Res. 40: D700–705

Cox J & Mann M (2008) MaxQuant enables high peptide identification rates, individualized p.p.b.-range mass accuracies and proteome-wide protein quantification. Nat Biotech 26: 1367–1372

Cox J, Neuhauser N, Michalski A, Scheltema RA, Olsen JV & Mann M (2011) Andromeda: A Peptide Search Engine Integrated into the MaxQuant Environment. J. Proteome Res. 10: 1794–1805

Dobin A, Davis CA, Schlesinger F, Drenkow J, Zaleski C, Jha S, Batut P, Chaisson M & Gingeras TR (2013) STAR: ultrafast universal RNA-seq aligner. Bioinformatics 29: 15–21

Fu C, Donovan WP, Shikapwashya-Hasser O, Ye X & Cole RH (2014) Hot Fusion: An Efficient Method to Clone Multiple DNA Fragments as Well as Inverted Repeats without Ligase. PLoS One 9: Available at: http://www.ncbi.nlm.nih.gov/pmc/articles/PMC4281135/ [Accessed September 11, 2017]

Garí E, Piedrafita L, Aldea M & Herrero E (1997) A set of vectors with a tetracycline-regulatable promoter system for modulated gene expression in Saccharomyces cerevisiae. Yeast 13: 837–848

Ghaemmaghami S, Huh W-K, Bower K, Howson RW, Belle A, Dephoure N, O’Shea EK & Weissman JS (2003) Global analysis of protein expression in yeast. Nature 425: 737–741

Giaever G, Chu AM, Ni L, Connelly C, Riles L, Véronneau S, Dow S, Lucau-Danila A, Anderson K, André B, Arkin AP, Astromoff A, El-Bakkoury M, Bangham R, Benito R, Brachat S, Campanaro S, Curtiss M, Davis K, Deutschbauer A, et al (2002) Functional profiling of the Saccharomyces cerevisiae genome. Nature 418: 387–391

Kushnirov VV (2000) Rapid and reliable protein extraction from yeast. Yeast 16: 857–860

Saveanu C, Rousselle J-C, Lenormand P, Namane A, Jacquier A & Fromont-Racine M (2007) The p21-activated protein kinase inhibitor Skb15 and its budding yeast homologue are 60S ribosome assembly factors. Mol. Cell. Biol. 27: 2897–2909

Wessel D & Flügge UI (1984) A method for the quantitative recovery of protein in dilute solution in the presence of detergents and lipids. Anal. Biochem. 138: 141–143

